# Distinct Proteasomal Pathways Drive Oncogenic PPM1D Activation

**DOI:** 10.64898/2026.06.01.729344

**Authors:** Ni Yan, Deumaya Shrestha, Cameron Schluter, Cyrus Jin, Edward Wang, Carmen Da Silva, Ilaria Gritti, Nikolaos Tsopoulidis, Ruxin Yang, Adam S. Sperling, Jonathan M. Tsai, Andrew E.H. Elia, Eric S. Fischer, Mikolaj Slabicki, Eugene Oh, Peter G. Miller

## Abstract

PPM1D is a serine/threonine phosphatase and DNA damage response (DDR) regulator recurrently activated in cancer through amplification or C-terminal truncating mutations that increase its abundance. Here we show that truncating mutations fundamentally rewire PPM1D proteostasis, unmasking an oncogenic function of an alternative protein degradation pathway. While full-length PPM1D undergoes rapid ubiquitin-independent proteasomal degradation via a C-terminal degron, truncating mutations redirect degradation to a slower UBR5-mediated ubiquitin-dependent pathway. The resulting accumulation of PPM1D suppresses DDR signaling and enhances cellular fitness under genotoxic stress, which is further amplified by UBR5 loss. Consistent with selective pressure on this axis, cancers harboring *PPM1D* truncating mutations are enriched for *UBR5* loss-of-function mutations. Together, these findings identify escape from ubiquitin-independent proteasomal degradation as a mechanism of oncogenic adaptation and establish proteostatic routing as a regulatory layer linking protein degradation, DDR signaling, and cancer evolution.

## INTRODUCTION

The protein phosphatase PPM1D (also known as WIP1) is a central regulator of the DNA damage response (DDR), acting through dephosphorylation of key signaling mediators including p53.^1–5^ Recurrent amplification, C-terminal truncating mutations, or increased transcription of *PPM1D* occur across a range of malignancies, most notably in therapy-related myeloid neoplasms, brain tumors, and breast cancers.^1,6^ In these oncogenic contexts, elevated PPM1D protein levels attenuate DDR signaling through dephosphorylation of checkpoint mediators, including p53 and other DDR substrates, enabling cancer cells and pre-malignant clones to tolerate genotoxic stress and acquire aggressive phenotypes such as therapy resistance.^7–12^ Accordingly, PPM1D has emerged as an attractive therapeutic target in multiple malignancies as its inhibition is predicted to restore DDR signaling and sensitize cells to cytotoxic therapy.

Previous studies, including our own, have shown that C-terminal truncating mutations in exon 6 produce a truncated protein (tr-PPM1D) with higher protein levels than full-length PPM1D (fl-PPM1D), but the mechanisms underlying this stabilization remain undefined.^10,11,13^ Accordingly, although the catalytic and regulatory functions of PPM1D have been extensively characterized, the mechanisms governing its protein turnover remain poorly defined. While the majority of proteasomal degradation relies on ubiquitination of target proteins mediated by E3 ligases in the ubiquitin-proteasome system (UPS), select substrates can be degraded through ubiquitin-independent proteasomal degradation (UbInPD).^14,15^ Whether PPM1D is subject to such noncanonical proteasomal regulation, and whether oncogenic truncating mutations alter its mode of proteasomal recognition, remain unknown. Moreover, whether proteasomal control of PPM1D abundance functions as a regulatory layer of DDR signaling remains unclear.

Here, we show that fl-PPM1D and tr-PPM1D are degraded by fundamentally distinct proteasomal mechanisms. fl-PPM1D undergoes ubiquitin-independent proteasomal degradation via its C-terminus, whereas truncating mutations eliminate this signal and shift PPM1D turnover to a UBR5-mediated, ubiquitin-dependent pathway. We further demonstrate that key previously described effects of UBR5 on DDR signaling are largely mediated through its regulation of tr-PPM1D. Consistent with selective pressure on this regulatory axis, cancers harboring *PPM1D* truncating mutations are enriched for *UBR5* loss-of-function alterations and lack co-occurring *UBR5* amplification. Together, these findings link the structural state of PPM1D to its mode of proteasomal regulation and define how recurrent oncogenic mutations rewire proteostatic control of DDR signaling.

## RESULTS

### Oncogenic Truncating Mutations Rewire PPM1D Proteasomal Degradation

Consistent with recurrent activation of PPM1D in human disease, amplifications, truncating mutations, and overexpression occur across cancer types, with a notable prevalence of truncating mutations in malignant and premalignant contexts including diffuse intrinsic pontine gliomas, therapy-related myeloid neoplasms, and clonal hematopoiesis (**Figures 1A-B and S1A**).^1,10,11,16–19^ We and others have shown that C-terminal frameshift or non-sense mutations in exon 6 of *PPM1D* result in expression of tr-PPM1D.^10,11,13,20,21^ We employed the MOLM13 acute myeloid leukemia cells, which is *PPM1D* wild-type and therefore expresses fl-PPM1D, to introduce indels into exon 6 of *PPM1D* at amino acid 495 via CRISPR/Cas9 to generate isogenic cells that express tr-PPM1D (**Figure 1C**). In parallel, we examined additional cell lines, including U2OS (p.R458*) and HCT116 (p.L450*), that harbor endogenous *PPM1D* truncating mutations and consequently express both tr-PPM1D and fl-PPM1D. Across all three systems, tr-PPM1D protein levels were higher than fl-PPM1D, and tr-PPM1D exhibited a longer half-life following treatment with cycloheximide, a drug that blocks protein translation and allows for quantification of protein stability and degradation (**Figures 1C-D**). Next, we treated cells with DMSO, the proteasome inhibitor MG132, or the autophagy inhibitor Bafilomycin A1 in the presence or absence of cycloheximide. Consistent with a proteasomal degradation, levels of both tr-PPM1D and fl-PPM1D increased at baseline and following cycloheximide treatment in the presence of MG132 but not Bafilomycin A1 (**Figure 1E**).

**Figure 1.**
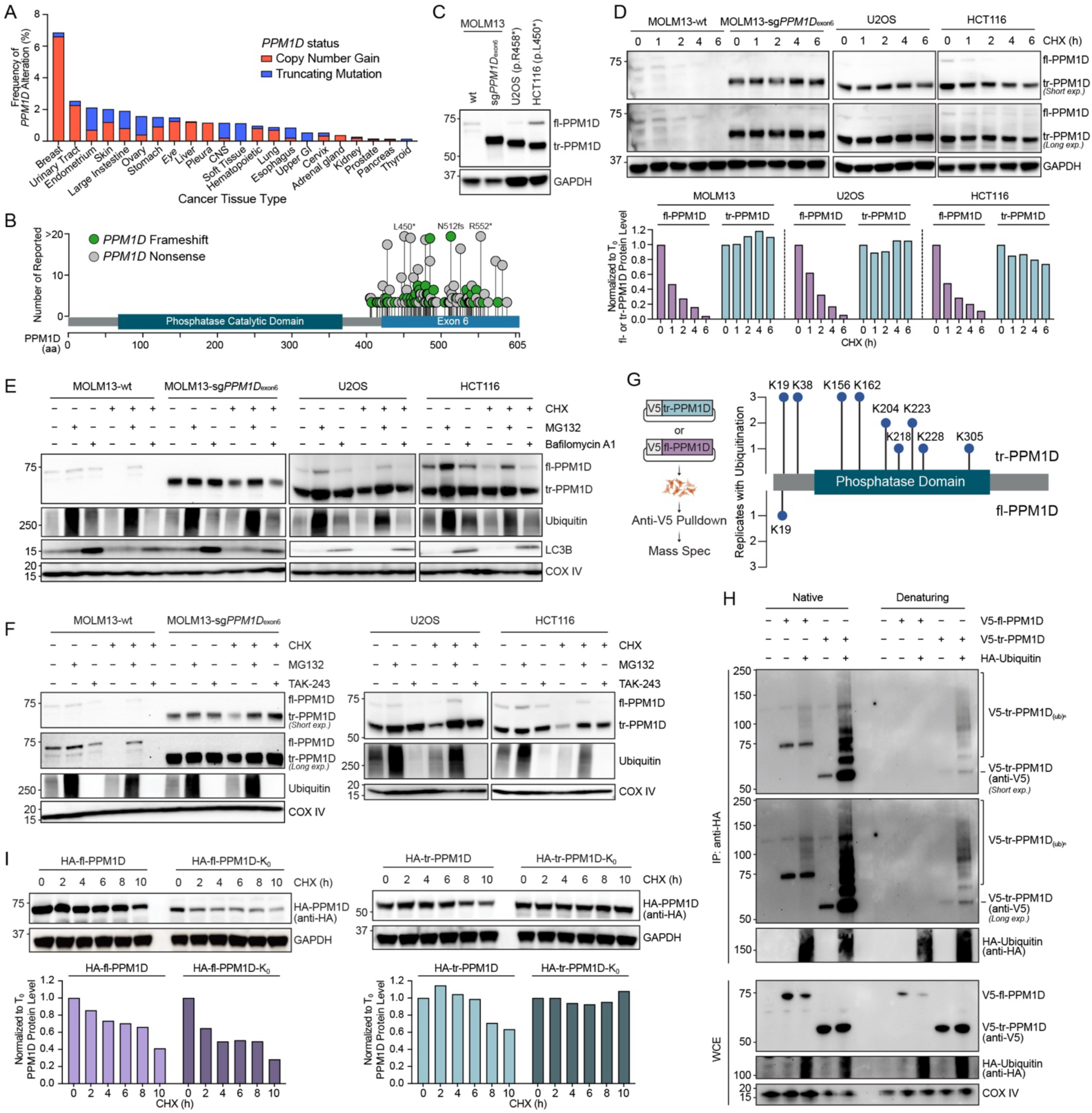
Oncogenic truncating mutations rewire PPM1D proteasomal degradation. (A) Histogram showing the frequency of *PPM1D* amplifications and C-terminal truncating mutations by cancer tissue type acquired from the Catalogue of Somatic Mutations in Cancer (COSMIC) database. Red box indicates *PPM1D* copy number gain, and blue box indicates *PPM1D* truncating mutation. (B) Lollipop plot showing the distribution of *PPM1D* C-terminal truncating mutations obtained from the COSMIC database. Green dot indicates *PPM1D* frameshift mutation, and gray dot indicates *PPM1D* nonsense mutation. Mutations with numbers of reported above 20 were labeled. (C) Western blot of PPM1D in MOLM13 wildtype (wt), MOLM13 truncated mutant (sg*PPM1D*_exon6_), U2OS, or HCT116 cells. (D) Western blot (top) and quantification (bottom) of a cycloheximide (CHX) chase time-course of PPM1D in MOLM13-wt, MOLM13-sg*PPM1D*_exon6_, U2OS, or HCT116. (E) Western blot of PPM1D, ubiquitin, and LC3B in MOLM13-wt, MOLM13-sg*PPM1D*_exon6_, U2OS, or HCT116 cells after treatment with CHX, MG132, or Bafilomycin A1. (F) Western blot of PPM1D and ubiquitin in MOLM13-wt, MOLM13-sg*PPM1D*_exon6_, U2OS, or HCT116 cells after treatment with CHX, MG132, or TAK-243. (G) Schematic of immunoprecipitation mass spectroscopy experiment where V5-tr-PPM1D or V5-fl-PPM1D were expressed in HEK293T cells (left) and quantification (right) of ubiquitinated lysines across replicates. (H) V5-fl-PPM1D or V5-tr-PPM1D were co-expressed with HA-Ubiquitin in HEK293T cells followed by native or denaturing immunoprecipitation of HA-Ubiquitin and immunoblotting for V5-PPM1D. Whole cell extract (WCE) is shown below. (I) Western blot (top) and quantification (bottom) of a CHX chase time-course of HA-PPM1D in HEK293T cells expressing HA-fl-PPM1D, HA-fl-PPM1D-K_0_, HA-tr-PPM1D, or HA-tr-PPM1D-K_0_.

To assess the role of the UPS, we treated cells with MG132 or TAK-243 (also known as MLN7243), a small-molecule inhibitor of the ubiquitin-activating enzyme UBA1 that globally suppresses ubiquitination.^22^ Whereas MG132 rescued degradation of both tr-PPM1D and fl-PPM1D, TAK-243 selectively rescued tr-PPM1D, indicating that tr-PPM1D degradation is ubiquitin-dependent (**Figure 1F**). In contrast, TAK-243 did not rescue fl-PPM1D degradation in any cell line tested, suggesting that fl-PPM1D undergoes UbInPD. Global suppression of ubiquitination with TAK-243 modestly accelerated fl-PPM1D turnover, as has been reported for other UbInPD substrates (**Figure S1B**).^23^

Next, we expressed V5-fl-PPM1D or V5-tr-PPM1D (PPM1D_1-426_) and performed immunoprecipitation mass spectrometry (IP-MS) to quantify the sites of protein ubiquitination. Despite identical N-terminal sequences, tr-PPM1D was ubiquitinated at many lysines across replicates, whereas fl-PPM1D ubiquitination was very sparse and inconsistent between replicates (**Figure 1G**). To confirm this finding, we co-expressed HA-Ubiquitin and V5-fl-PPM1D or V5-tr-PPM1D, immunoprecipitated HA-Ubiquitin under either native or denaturing lysis conditions and immunoblotted for V5. Under denaturing condition, only V5-tr-PPM1D exhibited laddering consistent with polyubiquitination (**Figure 1H)**. Using antibodies specific for K48- or K11-linked ubiquitin chains, which are chain types with high specificity for the proteasome, we found that tr-PPM1D ubiquitination was both K48- and K11-linked, consistent with proteasomal targeting (**Figure S1C**).^24^

To functionally test the role of ubiquitination in tr-PPM1D degradation, we expressed HA-tagged wild-type or a variant where all lysines were mutated to arginines (“K_0_”) of both PPM1D isoforms in HEK293T cells. Whereas HA-fl-PPM1D and HA-fl-PPM1D-K_0_ exhibited similar degradation kinetics after cycloheximide treatment and were unaffected by TAK-243, consistent with UbInPD, the tr-PPM1D-K_0_ mutant was stabilized and insensitive to TAK-243, highlighting the requirement for lysine ubiquitination in tr-PPM1D degradation (**Figures 1I and S1D**).

Together, these data demonstrate that oncogenic truncating mutations fundamentally alter PPM1D proteostasis by shifting degradation from a rapid, ubiquitin-independent mechanism to a slower, ubiquitin-dependent pathway, resulting in selective stabilization of tr-PPM1D.

### Ubiquitin-Independent Proteasomal Degradation of fl-PPM1D is Mediated by the C-terminus

Broadly, two mechanisms have been described for UbInPD: chaperone-mediated delivery, including midnolin (MIDN), ubiquilins, and heat-shock proteins, or direct engagement of the proteasome, as observed for substrates such as ornithine decarboxylase (ODC).^23,25,26^ However, knockdown of *MIDN* or *UBQLN1/2/4* in U2OS, HCT116, and HEK293T cells did not slow fl-PPM1D degradation, and overexpression of neither MIDN nor UBQLN1 enhanced fl-PPM1D degradation (**Figures S2A-B**).^23,27^ A recent report suggested that fl-PPM1D is targeted to the proteasome through PSMD14 and PSME3; however, we observed no significant effect of knockdown of *PSMD14*, *PSME3*, or the combination on fl-PPM1D degradation (**Figure S2C**).^28^ We also did not observe any effect of HSP70 or HSP90 inhibitors on fl-PPM1D degradation (**Figure S2D**).

To identify potential mediators of fl-PPM1D UbInPD, we performed a genome-wide CRISPR/Cas9 screen for regulators of the protein stability of fl-PPM1D and PPM1D_400-605_, which encodes the C-terminus of the protein. We generated lentiviral fluorescent reporters in which EGFP was fused in-frame to the N-terminus of tr-PPM1D, fl-PPM1D, or PPM1D_400-605_ and linked via an internal ribosome entry site (IRES) to mCherry for normalization.^29,30^ As expected, the tr-PPM1D reporter exhibited the highest GFP:mCherry ratio, consistent with increased stability, whereas MG132 treatment preferentially increased fl-PPM1D and PPM1D_400-605_ reporter levels, consistent with more rapid proteasomal degradation (**Figure 2A**). We next transduced the fl-PPM1D and PPM1D_400-605_ reporter cell lines with a genome-wide CRISPR/Cas9 knockout library. After ten days, cells were treated with DMSO or cycloheximide for six hours then sorted for the top 15% and bottom 15% of GFP:mCherry signal, corresponding to increased and decreased reporter protein levels, respectively (**Figure 2B**). Among the top genes whose loss stabilized both reporters in the presence of cycloheximide, we observed an enrichment for proteasomal components, with no clear identification of potential chaperone proteins (**Figures 2C-D, and Tables S1-2**).

**Figure 2.**
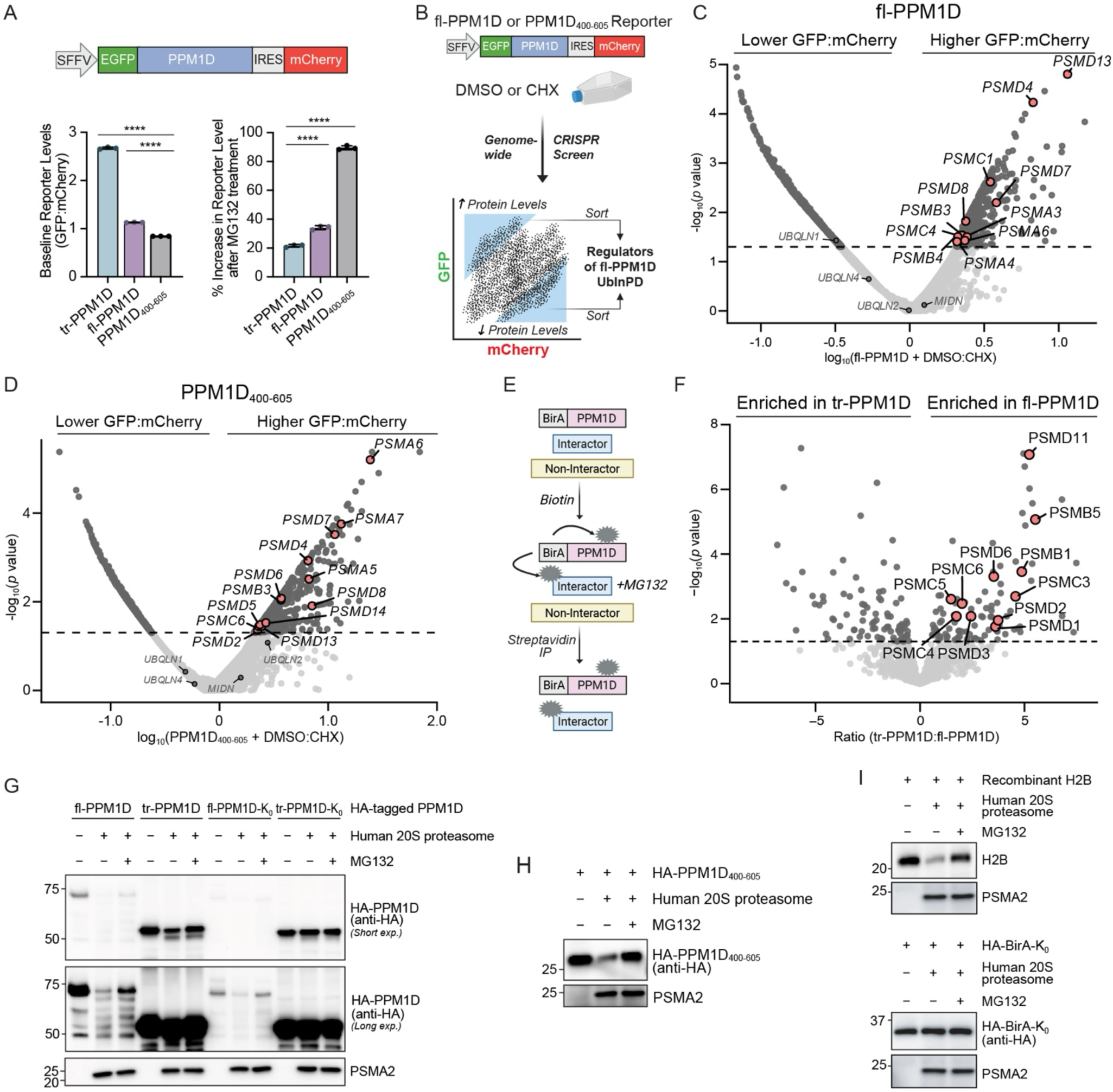
Ubiquitin-independent proteasomal degradation of fl-PPM1D. (A) GFP:mCherry quantification by flow cytometry in MOLM13 cells expressing the EGFP-tr-PPM1D-IRES-mCherry or EGFP-fl-PPM1D-IRES-mCherry or EGFP-PPM1D_400-605_-IRES-mCherry stability reporter at baseline (left) or after treatment with MG132 (right). (B) Schematic of a genome-wide CRISPR screen in U937 cells expressing the EGFP-fl-PPM1D-IRES-mCherry or EGFP-PPM1D_400-605_-IRES-mCherry stability reporter treated with DMSO or cycloheximide (CHX). (C) Volcano plot of a genome-wide CRISPR screen in U937 cells expressing the EGFP-fl-PPM1D-IRES-mCherry stability reporter treated with DMSO or CHX. (D) Volcano plot of a genome-wide CRISPR screen in U937 cells expressing the EGFP-PPM1D_400-605_-IRES-mCherry stability reporter treated with DMSO or CHX. (E) Schematic of a BirA-based proximity ligation assay in which HEK293T cells expressing BirA-tr-PPM1D or BirA-fl-PPM1D were incubated with biotin, followed by MG132 treatment. Biotinylated substrates were enriched by streptavidin immunoprecipitation and identified by mass spectrometry. (F) Volcano plot of a BirA-based proximity ligation assay in HEK293T cells expressing BirA-tr-PPM1D or BirA-fl-PPM1D. (G) Western blot of *in vitro* proteasome degradation assay. HA-tr-PPM1D, HA-tr-PPM1D-K_0_, HA-fl-PPM1D, or HA-fl-PPM1D-K_0_ was immunoprecipitated from HEK293T cells, the eluted PPM1D proteins were then incubated for 3 hours with recombinant human 20S proteasome with or without MG132. PSMA2 is a control to detect the proteasome. (H) Western blot of *in vitro* proteasome degradation assay. HA-PPM1D_400-605_ was immunoprecipitated from HEK293T cells, the eluted PPM1D protein was then incubated for 3 hours with recombinant human 20S proteasome with or without MG132. PSMA2 is a control to detect the proteasome. (I) Western blot of *in vitro* proteasome degradation assay. Recombinant Histone H2B (top) or HA-BirA-K_0_ immunoprecipitated from HEK293T cells (bottom) were incubated for 3 hours with recombinant human 20S proteasome with or without MG132. PSMA2 is a control to detect the proteasome.

To complement the CRISPR screen with an orthogonal proteomic approach, we performed proximity-labeling followed by mass spectrometry. We generated HEK293T cells expressing BirA fused in-frame to the N-terminus of fl-PPM1D and tr-PPM1D. We confirmed that the BirA-fusions did not change the mode of degradation of either fl-PPM1D or tr-PPM1D (**Figure S2E**). In the presence of biotin, BirA biotinylates nearby proteins, allowing for a Streptavidin pulldown (**Figure S2F**). Cells were treated with biotin and MG132, lysed, and subjected to streptavidin pulldown followed by mass spectrometry (**Figure 2E**). We observed a marked enrichment of proteasome-associated proteins in the fl-PPM1D condition compared with tr-PPM1D (**Figure 2F and Table S3**). Based on these data, we tested whether fl-PPM1D can be directly degraded by the proteasome. We immunoprecipitated HA-fl-PPM1D, HA-fl-PPM1D-K_0_, HA-tr-PPM1D, or HA-tr-PPM1D-K_0_ from HEK293T cells and incubated the eluates with purified 20S proteasome in the presence or absence of MG132. As expected, ubiquitinated tr-PPM1D was degraded, whereas tr-PPM1D-K_0_ was not, consistent with a ubiquitin-dependent degradation. In contrast, both fl-PPM1D and fl-PPM1D-K_0_ were degraded by 20S, consistent with direct, UbInPD (**Figure 2G**). We then confirmed that the C-terminus alone (amino acids 400-605) underwent *in vitro* degradation by 20S (**Figure 2H**). As important controls, we included H2B, a known direct substrate of 20S, and HA-BirA-K_0_ which is a highly stable protein not degraded by 20S (**Figure 2I**).^23,31^

We next investigated the sequence features that mediate UbInPD of fl-PPM1D. In our EGFP-fl-PPM1D-IRES-mCherry reporter, we generated a series of truncation mutations every 25 amino acids starting from the C-terminus. MOLM13 cells expressing these reporters were treated with DMSO or MG132, and reporter stability was quantified as the GFP:mCherry ratio under each condition. We observed that truncations beginning at amino acid 550 resulted in marked MG132-dependent stabilization, indicating that residues 550-605 encompass a degradation signal (**Figure 3A**). To confirm this finding, we expressed either HA-BirA-K_0_, a highly stable protein, or HA-BirA-K_0_ fused to PPM1D residues 550-605 (HA-BirA-K_0_-PPM1D_550-605_), treated cells with DMSO, MG132, or TAK-243, and assessed protein levels by immunoblotting.^23^ Whereas HA-BirA-K_0_ expression was high and unaffected by MG132 or TAK-243, HA-BirA-K_0_-PPM1D_550-605_ showed low baseline expression and was stabilized by MG132 but not TAK-243, a finding we validated using the GFP reporter system, confirming that residues 550-605 are sufficient for UbInPD (**Figures 3B and S3A**).

**Figure 3.**
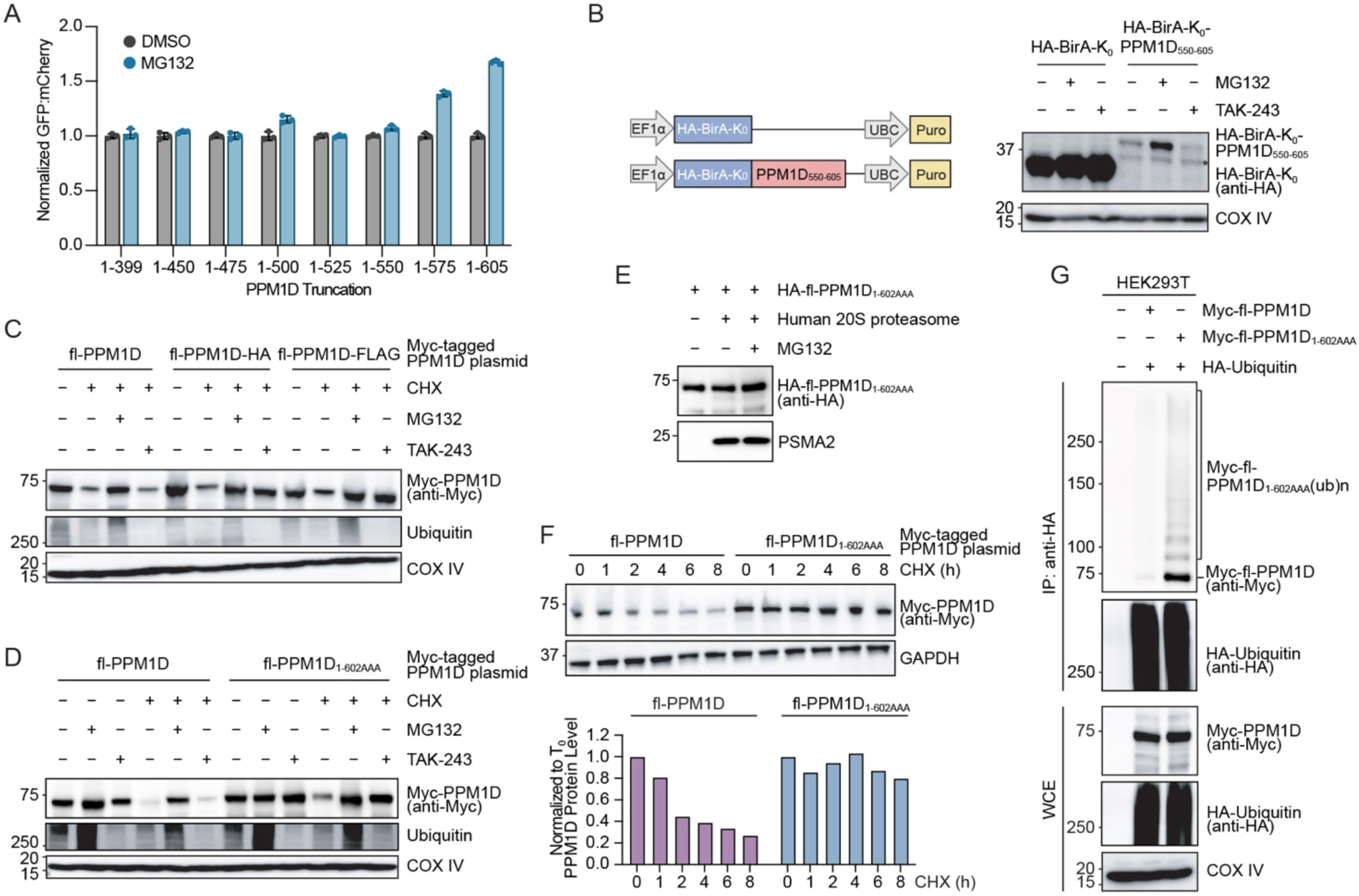
Ubiquitin-independent proteasomal degradation of fl-PPM1D is mediated by the C-terminus. (A) Normalized GFP:mCherry ratio detected by flow cytometry in MOLM13 cells expressing the EGFP-PPM1D-IRES-mCherry stability reporter with various C-terminal truncations treated with DMSO or MG132. Data are presented as mean ± SD (n = 3 biological replicates). (B) Schematic (left) and western blot (right) of HA-BirA-K_0_ in HEK293T cells expressing either HA-BirA-K_0_ or HA-BirA-K_0_-PPM1D_550-605_ after treatment with MG132 or TAK-243. Asterisk indicates an unspecific band from HA antibody. (C) Western blot of Myc-PPM1D in HEK293T cells expressing Myc-fl-PPM1D, Myc-fl-PPM1D-HA, or Myc-fl-PPM1D-FLAG after treatment with cycloheximide (CHX), MG132, or TAK-243. (D) Western blot of Myc-PPM1D in HEK293T cells expressing Myc-fl-PPM1D or Myc-fl-PPM1D_1-602AAA_ after treatment with CHX, MG132, or TAK-243. (E) Western blot of *in vitro* proteasome degradation assay. HA-fl-PPM1D_1-602AAA_ was immunoprecipitated from HEK293T cells, the eluted PPM1D protein was then incubated for 3 hours with recombinant human 20S proteasome with or without MG132. PSMA2 is a control to detect the proteasome. (F) Western blot (top) and quantification (bottom) of a CHX chase time-course of Myc-PPM1D in HEK293T cells expressing Myc-fl-PPM1D or Myc-fl-PPM1D_1-602AAA_. (G) Myc-fl-PPM1D or Myc-fl-PPM1D_1-602AAA_ were co-expressed with HA-Ubiquitin in HEK293T cells followed by denaturing immunoprecipitation of HA-ubiquitin and immunoblotting for Myc-PPM1D. Whole cell extract (WCE) is shown below.

To date, neither experimental nor computational approaches have defined the structure of the C-terminus of PPM1D, a region predicted to be highly disordered (**Figure S3B**).^32–34^ Because intrinsically disordered regions (IDRs) can be directly recognized by the proteasome, we hypothesized that disorder within the PPM1D C-terminus mediates fl-PPM1D degradation.^35^ However, this model was insufficient to explain fl-PPM1D degradation, as addition of HA or FLAG tag to the C-terminus converted fl-PPM1D to ubiquitin-dependent degradation, evidenced by TAK-243-mediated stabilization of fl-PPM1D-HA and fl-PPM1D-FLAG (**Figure 3C**). The three terminal residues in PPM1D are cysteine-valine-cysteine (CVC), and a recent report suggested that cysteine and valine residues are enriched in C-terminal degrons of substrates for UbInPD.^23^ Mutation of the terminal CVC motif to AAA converted fl-PPM1D to TAK-243-dependent degradation and rendered it resistant to 20S-mediated degradation in vitro, accompanied by increased stability and ubiquitination, implicating the CVC motif in UbInPD of fl-PPM1D (**Figures 3D-G**). However, addition of a C-terminal CVC motif to tr-PPM1D was insufficient to confer UbInPD (**Figure S3C**). Further, FANK1, the only other protein in the human proteome terminating in CVC, did not undergo UbInPD (**Figure S3D**), together indicating that the CVC motif alone is not sufficient to drive UbInPD. Importantly, neither tr-PPM1D nor FANK1 contain a C-terminal disordered region (**Figure S3E**), suggesting that both intrinsic disorder and the CVC motif may contribute to UbInPD of fl-PPM1D. Together, these data demonstrate that the C-terminus of fl-PPM1D engages the proteasome, leading to its rapid ubiquitin-independent turnover.

### UBR5 Mediates Ubiquitin-Dependent Degradation of Truncated PPM1D

Having defined the UbInPD of fl-PPM1D, we next sought to identify regulators of ubiquitin-dependent proteasomal degradation of tr-PPM1D using a UPS-targeted CRISPR screen (**Figure 4A and Table S4**). We transduced tr-PPM1D and fl-PPM1D reporter cell lines with a UPS-targeted library.^29^ After ten days, cells were sorted for the top and bottom 10% of GFP:mCherry signal, corresponding to increased or decreased reporter protein levels. This screen identified several candidate regulators, most notably UBR5 and OTUD5, whose loss selectively increased tr-PPM1D reporter levels without affecting fl-PPM1D, implicating UBR5 as a specific regulator of tr-PPM1D stability. UBR5 has been implicated in diverse cellular processes, including protein quality control, transcriptional regulation, and the DDR.^36–40^ Because OTUD5 regulates UBR5 stability, loss of OTUD5 phenocopies UBR5 loss.^41^

**Figure 4.**
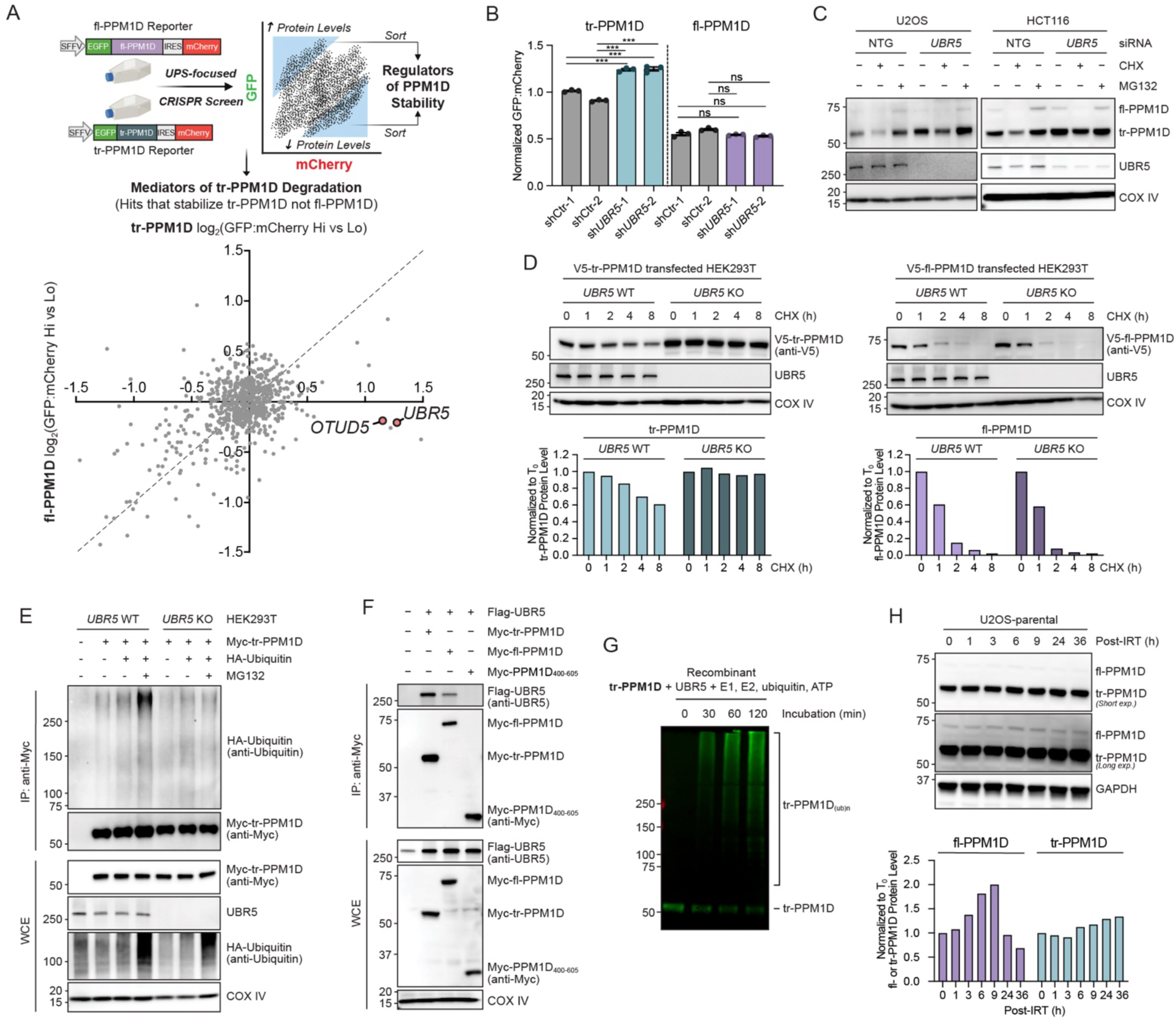
UBR5 mediates ubiquitin-dependent degradation of truncated PPM1D. (A) Schematic (top) and results (bottom) of a ubiquitin proteasome system (UPS) targeted CRISPR screen in U937 cells expressing the EGFP-fl-PPM1D-IRES-mCherry or EGFP-tr-PPM1D-IRES-mCherry stability reporter. (B) Normalized GFP:mCherry ratio detected by flow cytometry in U937 cells expressing the EGFP-fl-PPM1D-IRES-mCherry or EGFP-tr-PPM1D-IRES-mCherry stability reporter following control or *UBR5* knockdown using shRNAs. Data are presented as mean ± SEM (n = 3 biological replicates). Statistical significance was determined using unpaired two-tailed Student’s t-test; ns indicates non-significant (*p*>0.05), \*\*\**p*<0.001. (C) Western blot of PPM1D and UBR5 following siRNA mediated knockdown of *UBR5* and treatment with cycloheximide (CHX) or MG132 in U2OS or HCT116 cells. (D) Western blot (top) and quantification (bottom) of a CHX chase time-course of V5-PPM1D in *UBR5* wildtype (WT) or knockout (KO) HEK293T cells expressing either V5-tr-PPM1D (left) or V5-fl-PPM1D (right). (E) Myc-tr-PPM1D and HA-Ubiquitin were co-expressed in *UBR5* WT or KO HEK293T cells with or without MG132 treatment followed by denaturing immunoprecipitation of Myc-tr-PPM1D and immunoblotting for HA-Ubiquitin. Whole cell extract (WCE) is shown below. (F) Myc-tr-PPM1D, Myc-fl-PPM1D, or Myc-PPM1D_400-605_ were co-expressed with FLAG-UBR5 in HEK293T cells followed by immunoprecipitation of Myc-PPM1D and immunoblotting for UBR5. WCE is shown below. (G) Immunoblotting for tr-PPM1D following *in vitro* ubiquitylation assay performed by incubating recombinant tr-PPM1D and UBR5 with the necessary E1, E2, ubiquitin, and ATP substrates. (H) Western blot time-course (top) and quantification (bottom) of fl- or tr-PPM1D in U2OS parental cells following 2.5 Gy irradiation exposure.

Using shRNA, we confirmed that *UBR5* knockdown increased tr-PPM1D, but not fl-PPM1D, reporter levels in U937 cells (**Figure 4B**). Similarly*, UBR5* silencing selectively stabilized endogenous tr-PPM1D protein levels following cycloheximide treatment in U2OS and HCT116 cells without altering *PPM1D* mRNA levels (**Figures 4C and S4A**); this effect was phenocopied by *OTUD5* knockdown (**Figure S4B**). In *UBR5*-knockout HEK293T cells, tr-PPM1D, but not fl-PPM1D, was similarly stabilized following cycloheximide treatment (**Figure 4D**).^38^ Consistent with these findings, *UBR5* knockdown reduced tr-PPM1D ubiquitination, whereas UBR5 overexpression increased K48- and K11-linked ubiquitination (**Figures 4E and S4C**). We next co-expressed FLAG-UBR5 with Myc-tr-PPM1D, Myc-fl-PPM1D, or Myc-PPM1D_400-605_ in HEK293T cells. Following anti-Myc immunoprecipitation, UBR5 bound Myc-tr-PPM1D and Myc-fl-PPM1D, but not Myc-PPM1D_400-605_, indicating that UBR5 interacts with the N-terminus of PPM1D (**Figure 4F**). Finally, an *in vitro* ubiquitination assay using recombinant UBR5 demonstrated robust ubiquitination of tr-PPM1D, confirming that UBR5 directly ubiquitinates tr-PPM1D (**Figure 4G**).

DNA damage, including irradiation, drives transcription of PPM1D, resulting in a rise in protein levels then return to baseline as cells recover from the genotoxic insult.^3^ To quantify how differences in ubiquitin-independent and ubiquitin-dependent degradation pathways influence PPM1D kinetics, we performed a time-course analysis of fl-PPM1D and tr-PPM1D protein levels following 2.5 Gy irradiation. In parental U2OS cells, fl-PPM1D levels rose and declined to baseline within 24 hours, whereas tr-PPM1D levels, which were already significantly higher, increased further and remained elevated, consistent with slower degradation kinetics (**Figure 4H**). To interrogate this further, we used homology-directed repair to generate isogenic U2OS cell lines expressing either both tr-PPM1D and fl-PPM1D (control), fl-PPM1D alone, or tr-PPM1D alone at the endogenous locus (**Figure S4D**). We confirmed protein expression and appropriate degradation behavior in these isogenic cells: fl-PPM1D underwent UbInPD, whereas tr-PPM1D underwent UPS-mediated degradation (**Figure S4E**). Repeating the irradiation time course, we again observed a transient rise in fl-PPM1D protein levels, whereas tr-PPM1D protein levels rose and remained elevated even 36 hours after irradiation, consistent with prolonged suppression of DDR signaling reported in *PPM1D*-mutant cells (**Figure S4F**).^10,11,13,20,21^

Together, these data establish UBR5 as the E3 ligase that directly mediates ubiquitin-dependent proteasomal degradation of tr-PPM1D, thereby conferring slower degradation kinetics relative to fl-PPM1D.

### UBR5 Regulation of the DNA Damage Response Is Mediated by Truncated PPM1D

UBR5 has been implicated in diverse cellular processes, including protein quality control, transcriptional regulation, and modulation of the DDR.^36–40^ A prior study reported that UBR5 regulates the DDR through multiple mechanisms, including restriction of RNF168-mediated ubiquitination at sites of DNA damage and prevention of excessive ubiquitin spreading to undamaged chromatin.^40^ In this study, knockdown of *UBR5*, *TRIP12*, or both in U2OS cells enhanced DNA repair efficiency with a concomitant decrease in phosphorylation of KAP1 and γ-H2AX foci formation. Given the established role of PPM1D in DDR regulation, the presence of tr-PPM1D in U2OS cells, and our finding that UBR5 controls tr-PPM1D degradation, we hypothesized that the effects of UBR5 on these phenotypes are mediated by tr-PPM1D.

First, we knocked down *UBR5* or *PPM1D* by siRNA and induced DNA damage by irradiation across a panel of cell lines with defined PPM1D genotypes. In U2OS and HCT116 cells, which harbor tr-PPM1D, siRNA-mediated silencing of *UBR5* or the related E3 ligase *TRIP12* reduced p-KAP1 levels following 2.5 Gy irradiation, consistent with the prior report (**Figure S5A**).^40^ Importantly, this effect was abolished upon *PPM1D* silencing: in U2OS, HCT116, and BB49HNC cells (which also carry a heterozygous *PPM1D* C-terminal truncation), *UBR5* silencing no longer suppressed p-KAP1 levels when *PPM1D* was co-silenced, indicating that UBR5-dependent modulation of p-KAP1 levels is mediated through PPM1D (**Figures 5A and S5B**). To exclude off-target effects, we examined cell lines harboring both tr-PPM1D and loss-of-function *UBR5* mutations (SNUC5, CCK81, and SNU175). In these cells, *UBR5* silencing had no effect on tr-PPM1D abundance or on p-KAP1 levels following irradiation, confirming that the observed phenotype requires functional UBR5 (**Figure S5C**).

**Figure 5.**
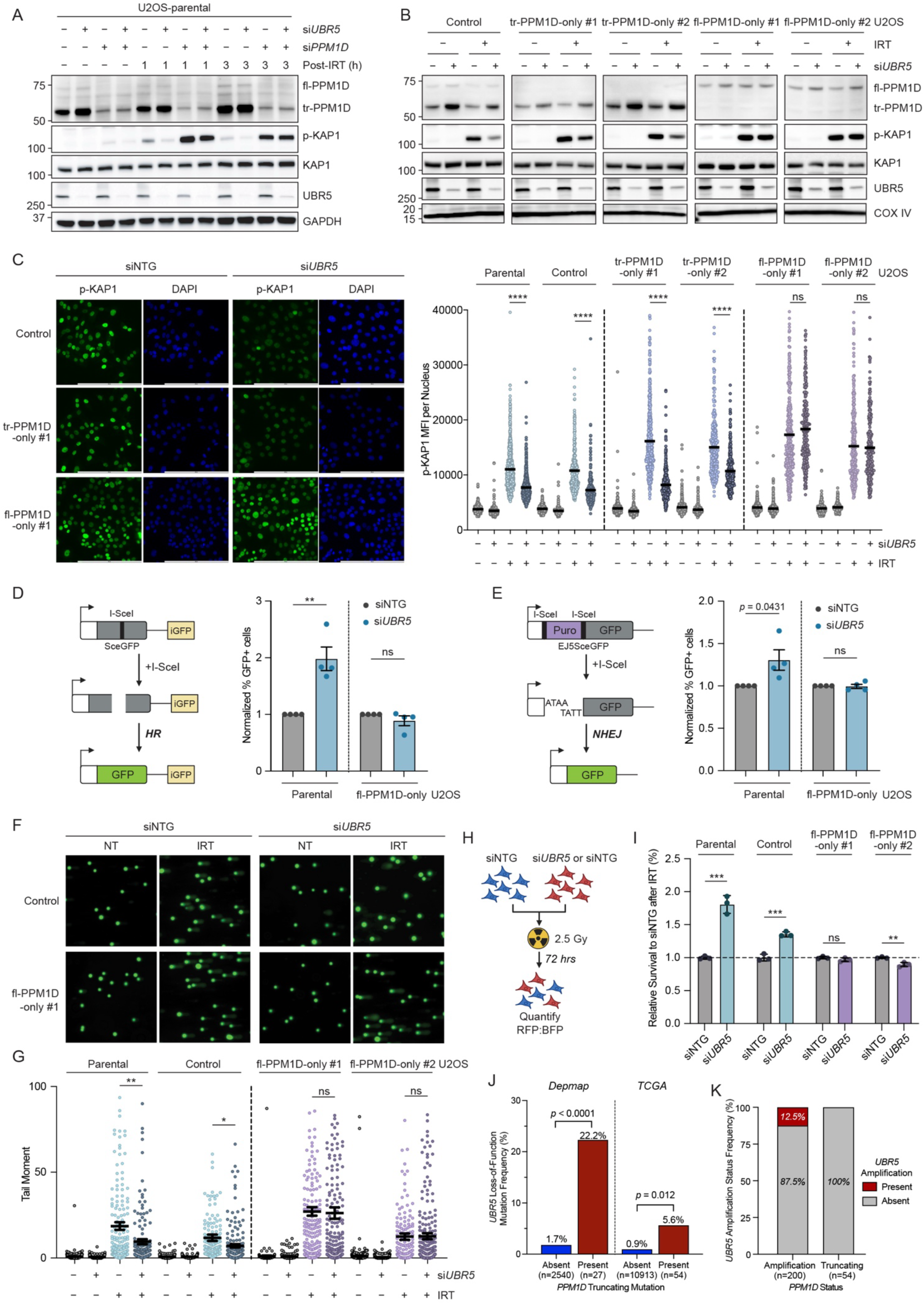
UBR5 regulation of the DNA damage response is mediated by truncated PPM1D. (A) Western blot time-course of PPM1D, p-KAP1, KAP1, and UBR5 in U2OS cells after siRNA mediated knockdown of non-targeting control (NTG), *UBR5*, *PPM1D,* or *UBR5* and *PPM1D* and exposure to 0 or 2.5 Gy irradiation. (B) Western blot of PPM1D, p-KAP1, KAP1, and UBR5 in U2OS isogenic cell lines after siRNA mediated knockdown of NTG or *UBR5* and exposure to 0 or 2.5 Gy irradiation. (C) Immunofluorescence analysis of p-KAP1 in U2OS parental and isogenic cell lines transfected with NTG or *UBR5* siRNA and exposed to 0 or 2.5 Gy irradiation. Representative images are shown on the left, and quantification of the mean fluorescence intensity (MFI) of p-KAP1 is shown on the right. n>230 cells for each condition were analyzed using CellProfiler. (D) Schematic of homology repair (HR) reporter (left) and normalized quantification (right) of GFP^+^ U2OS parental or fl-PPM1D-only isogenic cells following siRNA mediated knockdown of NTG or *UBR5*. (E) Schematic of non-homologous end joining (NHEJ) reporter (left) and normalized quantification (right) of GFP^+^ U2OS parental or fl-PPM1D-only isogenic cells following siRNA mediated knockdown of NTG or *UBR5*. (F) Representative images of COMET assay performed on U2OS parental or fl-PPM1D-only isogenic cells transfected with NTG or *UBR5* siRNA, 6 hours after treatment with 0 or 2.5 Gy irradiation. (G) Quantification of tail moments in COMET assay described in (F). n>160 comets for each condition were scored using OpenComet. (H) Schematic of cell growth competition assay. U2OS parental, control, or fl-PPM1D-only isogenic cells transfected with NTG or *UBR5* siRNA were fluorescently labeled with BFP or RFP, the cells were mixed, exposed to 2.5 Gy irradiation then grown for 72 hours prior to quantification of the RFP:BFP ratio. (I) Normalized quantification of RFP to BFP ratio in the competition assay described in (H). (J) *UBR5* truncating mutation frequency in *PPM1D* truncating mutation absent or present cell lines or primary tumors samples obtained from Depmap or TCGA. (K) *UBR5* amplification frequency in *PPM1D* amplification or truncating mutation patient samples obtained from TCGA. Data are presented as mean ± SEM, n = 4 biological replicates for Figures D and E, and n = 3 biological replicates for Figure I. For Figures D and E, each data point indicates the normalized mean of three internal biological replicates from each independent test. Statistical significance was determined using unpaired two-tailed Student’s t-test for Figures C, D, E, G, and I; using Fisher’s exact test for Figures J and K; ns indicates non-significant (*p*>0.05), \**p*<0.05, \*\**p*<0.01, \*\*\**p*<0.001, \*\*\*\**p*<0.0001.

To test whether this effect is specific to tr-PPM1D rather than fl-PPM1D, we utilized the U2OS isogenic cell lines. In the control and tr-PPM1D-only cells, *UBR5* knockdown decreased p-KAP1 levels after irradiation, whereas it had no effect on p-KAP1 levels in the fl-PPM1D-only cells, assessed by western blot, immunofluorescence, and intracellular flow cytometry (**Figures 5B-C and S5D**). We also examined cell lines that lack *PPM1D* truncations but overexpress fl-PPM1D through increased transcription (SKNDZ and NB1) or gene amplification (MCF7, KPL1, and IMR32).^42^ *UBR5* knockdown had no effect on p-KAP1 levels after irradiation in these cell lines, consistent with the findings from the U2OS isogenic models (**Figures S5E-F**). Together, these data indicate that UBR5-dependent regulation of p-KAP1 signaling is mediated by tr-PPM1D.

In the same prior study, *UBR5* knockdown in U2OS cells increased DNA repair, reflected by decreased γ-H2AX foci after irradiation and a concurrent increase in homologous recombination (HR) and non-homologous end joining (NHEJ).^40^ We therefore sought to determine whether these phenotypes were dependent on tr-PPM1D. We knocked down *UBR5* in the U2OS isogenic cell lines, treated the cells with or without 2.5 Gy irradiation then quantified γ-H2AX foci by immunofluorescence after 6 hours. Whereas γ-H2AX foci decreased after *UBR5* knockdown in parental and control cells, no significant change was observed in the fl-PPM1D-only cells (**Figure S6A**). Next, we used U2OS parental or fl-PPM1D-only cells expressing fluorescent reporters for HR (DR-GFP) or NHEJ (EJ5-GFP), in which I-SceI transfection generates a defined double-strand break that restores GFP expression upon repair by HR or NHEJ.^43,44^ In parental U2OS cells, *UBR5* knockdown increased the fraction of GFP^+^ cells in both reporters, particularly HR, whereas *UBR5* knockdown in fl-PPM1D-only cells had no effect on either mode of DNA repair (**Figures 5D-E**). We then directly quantified DNA breaks using the alkaline COMET assay in the isogenic cells. *UBR5* knockdown decreased the COMET tail moments after irradiation in both parental and control cells, whereas there was no change in the fl-PPM1D-only cells (**Figures 5F-G**). Taken together, these data indicate that UBR5-dependent regulation of DNA repair in U2OS cells is mediated by tr-PPM1D rather than fl-PPM1D.

Given that *UBR5* silencing enhanced DNA repair and tolerance to genotoxic stress in a tr-PPM1D-dependent manner, we next asked whether the UBR5-tr-PPM1D axis influences cellular sensitivity to radiation. To do so, we performed growth competition assays by introducing fluorescent markers into the parental, control, and fl-PPM1D U2OS cell lines (**Figure 5H**). In agreement with the prior report, *UBR5* knockdown conferred a competitive advantage in parental and control U2OS cells. However, this effect was eliminated in the fl-PPM1D-only cells (**Figure 5I**), consistent with a requirement for tr-PPM1D to mediate the fitness effects of *UBR5* silencing after irradiation.

Finally, having shown that the UBR5-tr-PPM1D axis regulates DDR and cellular fitness, we hypothesized that cancers harboring *PPM1D* truncating mutations would be under selective pressure to lose UBR5 function to further enhance tr-PPM1D levels. Although uncommon, *UBR5* loss-of-function mutations occur across multiple cancer types, typically through disruption of the C-terminal HECT domain.^45^ To test this hypothesis, we analyzed 2,567 cell lines from the Cancer Dependency Map and identified predicted loss-of-function *UBR5* mutations (restricted to frameshift mutations to enrich for impaired HECT domain function) in 22% of *PPM1D*-truncated lines (6/27) compared to 1.7% of cell lines lacking *PPM1D* truncating mutations (43/2540) (*p*<0.0001, Fisher’s exact test). This enrichment was also observed in clinical samples: among 10,967 TCGA tumors, *UBR5* mutations were present in 5.6% of *PPM1D*-truncated tumors (3/54) versus 0.9% of tumors without *PPM1D* truncating mutations (93/10913) (*p*=0.012, Fisher’s exact test) (**Figure 5J**). Conversely, *UBR5* amplifications (defined by GISTIC) were absent in *PPM1D*-truncated cases, whereas they were observed in tumors with full-length *PPM1D* amplification (0/54 vs. 25/200)), indicating that this pattern is specific to truncating mutations rather than PPM1D activation more broadly (**Figure 5K**).^46^ Together, these functional and genetic data link UBR5-dependent control of tr-PPM1D stability to DDR regulation and cellular fitness under genotoxic stress, highlighting allele-specific selective pressure on a degradation-dependent regulatory axis in cancers with *PPM1D* truncating mutations.

## DISCUSSION

PPM1D is a recurrently activated oncogene and central regulator of the DDR; however despite recurrent amplifications and truncating mutations observed in cancer, the mechanisms governing its protein stability have remained poorly defined.^1^ Here, we demonstrate that fl-PPM1D and tr-PPM1D are degraded by fundamentally distinct proteasomal pathways. Whereas tr-PPM1D undergoes ubiquitin-dependent degradation mediated by the E3 ligase UBR5, fl-PPM1D is subject to rapid ubiquitin-independent proteasomal degradation through engagement of the proteasome via a C-terminal degron containing a CVC motif. These findings provide a mechanistic explanation for the selective stabilization of tr-PPM1D in cancer.

Our data support a model in which fl-PPM1D is normally short-lived due to direct proteasomal recognition of its C-terminal degron. This model explains how cancer cells can overcome this constraint through *PPM1D* amplification or increased transcription, whereas truncating mutations that remove the degron more effectively bypass this rapid degradation pathway. The resulting tr-PPM1D retains catalytic activity and suppresses DDR signaling, including p53 activation.^12,21,33^ In this context, UBR5 provides a secondary quality-control mechanism by ubiquitinating tr-PPM1D leading to its degradation, albeit with far lower efficiently than the direct proteasomal route governing the full-length protein.^38^ The rarity of germline *PPM1D* truncations suggests that no dedicated surveillance mechanism has evolved for this mutant protein, leaving cells reliant on broader ubiquitin-dependent pathways.^47^

These findings place PPM1D among a limited class of proteins subject to UbInPD.^15,23,26,27,48^ The C-terminal region of fl-PPM1D, which is predicted to be intrinsically disordered and contains a terminal CVC motif, appears to function as a composite degron that contributes to proteasomal engagement. Disruption of this region through truncation, epitope tagging, or mutation of the CVC motif converts degradation to a ubiquitin-dependent mode, suggesting that both intrinsic disorder and sequence-specific features contribute to proteasomal routing. Notably, these findings suggest that UbInPD can function as a regulatory mechanism for DDR signaling through constitutive degradation of PPM1D. By maintaining low steady-state levels of fl-PPM1D, UbInPD limits basal PPM1D protein levels and thereby restrains phosphatase-mediated attenuation of DDR signaling, whereas escape from this pathway via truncating mutations enables sustained attenuation of checkpoint signaling.

Identification of UBR5 as the E3 ligase controlling tr-PPM1D turnover also clarifies prior observations linking UBR5 to DDR regulation. UBR5 influences chromatin ubiquitination and DDR marker accumulation through regulation of RNF168 protein levels.^40^ Our data demonstrate that UBR5-dependent attenuation of DDR signaling, including reduced p-KAP1 phosphorylation, decreased γ-H2AX foci, enhanced DNA damage repair, and improved tolerance to genotoxic stress, is dependent on tr-PPM1D, consistent with prior studies linking PPM1D activity with enhanced DNA repair.^49–51^ These phenotypes are abolished upon PPM1D silencing or in cells expressing only fl-PPM1D, establishing functional epistasis between UBR5 and mutant PPM1D. Loss of UBR5 stabilizes tr-PPM1D leading to enhanced DNA repair and decreased sensitivity to genotoxic stress. Consistent with these findings, *UBR5* loss-of-function mutations are enriched in *PPM1D*-mutant cancer cell lines and primary tumors, supporting selective pressure to further amplify tr-PPM1D signaling in human malignancies.

Beyond its mechanistic implications, these findings have direct relevance for the role of *PPM1D* truncating mutations in cancer. These mutations are recurrent across both hematologic and solid malignancies as well as clonal hematopoiesis, where they confer increased protein stability and attenuate DDR signaling through sustained dephosphorylation of checkpoint substrates including p53.^1^ This dampened DDR is thought to promote tolerance to genotoxic stress, enabling continued proliferation under conditions of DNA damage and contributing to genomic instability and therapy resistance. Our data extend this model by demonstrating that stabilization of tr-PPM1D through loss of *UBR5* further enhances DNA damage tolerance and cellular fitness, suggesting that the UBR5-tr-PPM1D axis functions as a proteostatic constraint on oncogenic PPM1D activity in cancer cells.

Together, these findings reveal that recurrent oncogenic truncating mutations in *PPM1D* rewire proteostatic routing by shifting PPM1D from rapid ubiquitin-independent proteasomal degradation to a slower ubiquitin-dependent pathway, thereby suppressing DDR signaling and enhancing cellular fitness. Loss of UBR5 further stabilizes tr-PPM1D, increasing mutant protein levels and reinforcing suppression of DDR signaling. In this context, cancers harboring *PPM1D* truncating mutations may be particularly dependent on mechanisms that restrain tr-PPM1D stability following escape from UbInPD, highlighting proteostatic control of mutant PPM1D as a potential therapeutic vulnerability. More broadly, this work altered proteostatic routing of PPM1D as a mechanism of oncogenic adaptation and suggests that restoring regulated PPM1D turnover may represent a strategy to re-engage DDR signaling in malignant and pre-malignant contexts.

## LIMITATIONS OF STUDY

While our findings uncover distinct proteasomal mechanisms controlling fl-PPM1D and tr-PPM1D, several limitations warrant consideration. Most experiments were performed in transformed cell lines, which may not fully recapitulate the proteostatic environment of primary hematopoietic or tumor cells; consequently, the contribution of additional accessory cofactors to UBR5- or proteasome-mediated degradation remains to be defined. The biochemical interaction between the PPM1D C-terminal degron and the 20S proteasome, while supported by our data, will require structural resolution to define the molecular recognition interface, as the structure of the C-terminus remains undefined. Similarly, the upstream cues that regulate UBR5 engagement of tr-PPM1D, including post-translational modifications or partner interactions, remain unknown. Finally, while we demonstrate that UBR5 modulates DDR signaling through tr-PPM1D, the broader impact of this pathway on genome stability, clonal selection, and therapeutic response will require validation in physiologic models.

## RESOURCE AVAILABILITY

### Lead Contact

Further information and requests for resources should be directed to and will be fulfilled by the lead contact, Peter Miller.

### Materials availability

Requests for generated materials should be directed to the corresponding authors.

### Data and code availability

Requests for data should be directed to the corresponding authors.

## ACKNOWLEDGMENTS

P.G.M. was supported by the National Cancer Institute (K08-CA263181) and the Edward P. Evans Foundation. This work was further supported by the American Cancer Society. We thank Dr. Peggy Goodell for providing the HR and NHEJ reporter cell lines, Dr. Michael Rape for providing the *UBR5* knockout cell lines, Dr. Ross Tomaino for support with proteomics experiments, and Dr. Christopher Ott for providing cell lines. We also thank Dr. Abby Green and Dr. Josephine Kahn for their thoughtful review of the manuscript.

## AUTHOR CONTRIBUTIONS

N.Y., D.S., C.S., C.J., C.D.S, I.G, N.T, R.Y. performed the experiments. N.Y., M.S., C.J., J.T., A.S.S., E.O., and P.G.M, designed the experiments. N.Y., D.S., C.J., E.W., A.E., E.S.F., M.S., and P.G.M. analyzed the data. N.Y. and P.G.M. wrote the paper. All authors read and accepted the manuscript. N.Y. and P.G.M. conceived the study. P.G.M. supervised the research.

## EXPERIMENTAL MODEL AND STUDY PARTICIPANT DETAILS

### Mammalian cell culture

MOLM13 and U937 cells are commercially available and were maintained in RPMI 1640 Medium (Thermo Fisher Scientific, 11875093) supplemented with 10% fetal bovine serum (FBS) (Omega Scientific, FB-02) and 1× penicillin-streptomycin (Thermo Fisher Scientific, 15140122). U2OS, HCT116, HEK293T, MCF7, BB49HNC, SNUC5, CCK81, SNU175, KPL1, IMR32, SKNDZ, and NB1 cells are commercially available and were maintained in Dulbecco’s Modified Eagle’s Medium (DMEM) (Sigma-Aldrich, D6429) supplemented with 10% FBS, 1× penicillin-streptomycin, and 1× L-Glutamine (Thermo Fisher Scientific, 25030081). The *UBR5* knockout and control HEK293T cell lines are a gift from Dr. Michael Rape’s lab. The U2OS parental and fl-PPM1D-only cells expressing fluorescent reporters for homology repair (HR) (DR-GFP) or non-homologous end joining (NHEJ) (EJ5-GFP) reporters were kindly provided by Dr. Peggy Goodell’s lab. U2OS-control, U2OS-tr-PPM1D-only, and U2OS-fl-PPM1D-only cells were generated by CRISPR/Cas9 editing in our lab. All cell lines were maintained at 37 °C and 5% CO_2_.

All cell lines were routinely tested for mycoplasma contamination using the Mycoplasma Detection Kit (Vazyme, D201).

## METHOD DETAILS

### Plasmids

Plasmids expressing V5-tagged fl-PPM1D, tr-PPM1D, or PPM1D_400-605_ were generated as previously described.^1^ Plasmids expressing V5-, HA- or Myc-tagged fl-PPM1D or tr-PPM1D, fl-PPM1D-HA or -FLAG, fl-PPM1D_1-602AAA_, or tr-PPM1D-CVC were generated based on V5-PPM1D plasmids using the Q5 Site-Directed Mutagenesis Kit (NEB, E0554S). EGFP-PPM1D-IRES-mCherry reporters, HA-PPM1D-K_0_, BirA-PPM1D, HA-BirA-K_0_, HA-BirA-K_0_-PPM1D_550-605_, Flag-Luciferase, Flag-MIDN, Flag-UBQLN1, and Myc-FANK1 plasmids were synthesized by Twist Bioscience. HA-Ubiquitin (Addgene, 17608) plasmid was obtained from Addgene. The FLAG-tagged UBR5 plasmid was a gift from Dr. Jonathan Tsai.

### Transfection and transduction experiments Plasmid transfection experiments

Plasmid transfections for overexpression in HEK293T cells were performed using the TransIT-LT1 transfection reagent (Mirus Bio, MIR 2304) according to manufacturer’s instruction. Cells were processed for downstream experiments 48 hours after plasmids transfection.

### Lentivirus transduction experiments

Lentiviruses were produced in HEK293T cells by co-transfection of lentiviral and packaging plasmids using TransIT-LT1 transfection reagent. Viruses were harvested 48 hours post transfection, aliquoted, and stored at - 80 °C for later use.

One million suspension cells per well were seeded in a 6-well plate and resuspended in 0.5 mL of culture medium. A total of 2 mL of viral supernatant and polybrene (final concentration, 6 µg/mL) were added to each well. Plates were centrifuged at 2,250 rpm for 2 hours at 37 °C. Following centrifugation, 2.5 mL of fresh culture medium was added to each well, and the plates were returned to the incubator.

### siRNA knockdown

siRNA transfections for knockdown specific genes were performed using 15 pmol indicated siRNA and 1.5 µL Lipofectamine 3000 Reagent (Thermo Fisher Scientific, L3000008) per well in 24-well plate. Transfections in 6-well plates were scaled up according to the manufacturer’s instructions. Cells were treated 72 hours after siRNA transfection and subsequently collected for analysis. Non-targeting control siRNA pools (D-001206-13-05), siRNAs targeting *UBR5* (M-007189-02-0005), *MIDN* (M-023894-01-0005), *UBQLN1* (M-012942-01-0005), *UBQLN2* (M-013566-00-0005), *UBQLN4* (M-021178-00-0005), *PSMD14* (M-006024-00-0005), *PSME3* (M-012133-00-0005 ), *OTUD5* (M-013823-00-0005), *TRIP12* (M-007182-01-0005), and *PPM1D* (M-004554-00-0005) were purchased from Dharmacon.

### Small molecule inhibitor treatment

For degradation assay, cells were pre-treated with DMSO, 10 µM MG132 (Selleck, S2619), or 1 µM TAK-243 (MedChemExpress, HY-100487) for 1 hour, then with or without 50 µg/mL cycloheximide (Selleck, S7418) for the indicated time. For HSP90 and HSP70 inhibition assay, cells were treated with 0.5 µM SNX-2112 (Selleck, S2639), 1 µM BP3 (MedChemExpress, HY-115997), 0.5 µM Ganetespib (Selleck, S1159), 1 µM Tanespimycin (Selleck, S1141), 1 µM BIIB021 (Selleck, S1175), or 20 µM VER-155008 (Selleck, S7751) for 24 hours.

### Generation of CRISPR/Cas9 HDR cell lines

Alt-R sgRNAs targeting *PPM1D*, Alt-R HDR donor oligo to reverse the truncated *PPM1D* allele to full-length *PPM1D*, and Cas9-GFP nuclease (IDT, 10008100) were introduced to U2OS parental cells according to manufacturer’s protocol using Cell Line Nucleofector Kit V (Lonza, VCA-1003) on a Nucleofector 2b device. Twenty-four hours after nucleofection, the culture medium was gently replaced with fresh medium. Seventy-two hours after nucleofection, GFP-positive cells were single-cell sorted into 96-well plates. Clonal populations were expanded, and *PPM1D* editing was verified by immunoblotting. Oligos sequences can be found in Table S5.

### Immunoprecipitation (IP)

For in-cell ubiquitin IP, HEK293T cells were transfected with V5- or Myc-tagged PPM1D and HA-Ubiquitin plasmids. Forty-eight hours after transfection, cells were treated with DMSO, 5 µM MG132, or 5 µM MG132 and 5 µM cycloheximide for 3 hours. Cells were then collected and lysed in NP40 buffer (50 mM Tris-HCl, pH 8, 150 mM NaCl, 1 mM EDTA, 1% NP40) supplemented with 10 µM MG132, 1× Halt Protease Inhibitor Cocktail (Thermo Fisher Scientific, 78438) and 10 mM N-Ethylmaleimide (MedChemExpress, HY-D0843). For denaturing condition, sodium dodecyl sulfate (SDS) was added to the lysates to a final concentration of 1%, followed by boiling at 95 °C for 6 minutes. After cooling to room temperature, lysates were diluted with IP buffer to reduce the SDS concentration to below 0.1%. Samples were then sonicated at 4 °C and clarified by centrifugation at 14,000 rpm for 30 minutes at 4 °C. For each sample, 30 µL of pre-equilibrated V5- (Proteintech, v5tma), Myc-(AlpaLifeBio, KTSM1336), or HA- (AlpaLifeBio, KTSM1335) magnetic agarose beads slurry was added. Lysate–bead mixtures were incubated at 4 °C for 3 hours with end-to-end rotation. Beads were subsequently washed six times with IP buffer, and bound proteins were eluted by boiling in 2× SDS sample buffer. Eluted proteins were analyzed by immunoblotting.

To assess the interaction between UBR5 and PPM1D, HEK293T cells were transfected with FLAG-UBR5 and Myc-PPM1D plasmids. Forty-eight hours after transfection, cells were harvested and lysed in NP40 buffer supplemented with 1× Halt Protease Inhibitor Cocktail. Lysates were sonicated, clarified by centrifugation, and immunoprecipitation was performed following the same procedure described above.

### Immunoblotting

Cells were lysed in SDS lysis buffer (NP40 buffer supplemented with 1% SDS) containing 1× Halt Protease Inhibitor Cocktail and 1× Halt Phosphatase Inhibitor (Thermo Fisher Scientific, 78420), followed by boiling at 100 °C for 10 minutes. Protein concentrations were determined using the Pierce BCA Protein Assay Kit (Thermo Fisher Scientific, 23227). Ten to twenty micrograms of total protein were resolved by SDS–PAGE (GenScript, M00657) and transferred onto 0.45 µm PVDF membranes (Sigma-Aldrich, IPFL00010). Membranes were blocked with 3% bovine serum albumin (BSA) (Sigma-Aldrich, A7906) in Tris-buffered saline containing 0.1% Tween-20 (TBST) for 1 hour at room temperature and incubated overnight at 4 °C with primary antibodies diluted in 3% BSA in TBST. After washing, membranes were incubated with HRP-conjugated secondary antibodies for 1 hour at room temperature. Immunoreactive signals were detected using enhanced chemiluminescence (ECL) reagents (Thermo Fisher Scientific, 34580). The following antibodies for immunoblotting were used in this study: PPM1D (Cell Signaling Technology, 11901), PPM1D (Santa Cruz Biotechnology, sc-376257), GAPDH (Cell Signaling Technology, 2118), Ubiquitin (Santa Cruz Biotechnology, sc-8017), LC3B (Cell Signaling Technology, 2775), COX IV (Cell Signaling Technology, 4844), HA (Cell Signaling Technology, 3724), V5 (Abcam, ab309485), Myc (Cell Signaling Technology, 2272), PSMA2 (Cell Signaling Technology, 2455), UBR5 (Cell Signaling Technology, 65344), p-KAP1 (S824) (Bethyl Laboratories, A300-767A), KAP1 (Cell Signaling Technology, 4124), K11-linked ubiquitin (kindly provided by Dr. Eugene Oh), K48-linked ubiquitin (kindly provided by Dr. Eugene Oh), FLAG (Thermo Fisher Scientific, MA1-91878), PSMD14 (Cell Signaling Technology, 4197), PSME3 (Proteintech, 14907-1-AP), H2B (Cell Signaling Technology, 12364), OTUD5 (Cell Signaling Technology, 20087), TRIP12 (Proteintech, 25303-1-AP), HRP-conjugated Affinipure Goat Anti-Rabbit IgG(H+L) (Proteintech, SA00001-2), and HRP-conjugated Affinipure Goat Anti-Mouse IgG(H+L) (Proteintech, SA00001-1).

### Immunofluorescence (IF)

U2OS cells were seeded in 6-well plates at approximately 60% confluency. Forty-eight hours after transfection of *UBR5* siRNA (Dharmacon, M-007189-02-0005) or non-targeting control siRNA (Dharmacon, D-001206-13-05), cells were re-seeded onto poly-L-lysine–coated glass coverslips (Sigma-Aldrich, P8920) placed in 6-well plates. The following day, cells were exposed to 2.5 Gy of irradiation. For detection of phospho-KAP1 (S824) (Bethyl Laboratories, A300-767A), IF was performed 1 hour after irradiation, whereas IF for γ-H2AX (S139) (BioLegend, 613402) was performed 6 hours after irradiation. Cells were fixed with 4% paraformaldehyde (PFA) (Santa Cruz Biotechnology, sc-281692) for 20 minutes, permeabilized with 0.1% Triton X-100 (Sigma-Aldrich, T9284) for 30 minutes, and blocked with 1% BSA/PBS for 1 hour at room temperature. Cells were then incubated overnight at 4 °C with primary antibodies diluted in 1% BSA/PBS. After washing, cells were incubated with anti-rabbit Alexa Fluor 488 (Thermo Fisher Scientific, A-21206), or anti-mouse Alexa Fluor 647 (Thermo Fisher Scientific, A-21236) secondary antibodies for 1 hour at room temperature protected from light. Coverslips were mounted using DAPI-containing mounting medium (Vector Laboratories, H-1200-10). Images were acquired using a Nikon Eclipse Ti inverted microscope.

### Intracellular flow cytometry for p-KAP1 and KAP1

U2OS parental and isogenic cell lines were transfected with *UBR5* or non-targeting control siRNA. Three days after transfection, cells were exposed to 2.5 Gy irradiation. One hour after irradiation, cells were trypsinized and stained with a fixable viability dye (Thermo Fisher Scientific, 65-0866). Cells were then fixed with 4% PFA in PBS at room temperature for 20 min, followed by permeabilization with ice-cold 90% methanol on ice for 30 min. After washing, cells were blocked with Fc-block (BD Pharmingen, 553141) in 1% BSA/PBS at 4 °C for 10 min and then incubated with phospho-KAP1 (S824) antibody (Bethyl Laboratories, A300-767A) diluted in 1% BSA/PBS at room temperature for 1 hour. Cells were washed and incubated with anti-rabbit Alexa Fluor 488 secondary antibody (Thermo Fisher Scientific, A-11034) and Alexa Fluor 647–conjugated KAP1 antibody (BioLegend, 619308) diluted in 1% BSA/PBS at room temperature for 1 hour protected from light. After final washes, cells were analyzed by flow cytometry.

### Real-time qPCR (RT-qPCR)

For qRT-PCR analysis, total RNA was purified form cells using the FastPure Cell/Tissue Total RNA Isolation Kit V2 (Vazyme, RC112). For each sample, 1 µg of total RNA was reverse transcribed using the HiScript III 1st Strand cDNA Synthesis Kit (Vazyme, R312) and then diluted 10-fold for RT-qPCR. Expression levels were quantified using Taq Pro Universal SYBR qPCR Master Mix (Vazyme, Q712) on a Bio-Rad CFX384 Real Time PCR System. RT-qPCR primers used in this study can be found in Table S5.

### Protein expression and purification

3xFLAG-tagged UBR5 was transfected in Expi293 cells (Thermo Fisher Scientific, A14635) following the manufacturer’s protocol. Cells were harvested 60 hours post-transfection and lysed in 50 mM HEPES pH 7.5, 200 mM NaCl, 0.5 mM TCEP supplemented with protease inhibitors. Lysate was cleared by ultracentrifugation (120,000g, 60 minutes) and incubated with anti-Flag resin for 1 hour. Resin was washed with lysis buffer and eluted with 0.2 mg/mL 3xFLAG peptide. Elution containing UBR5 was concentrated and further polished on Superose6Increase in buffer containing 25 mM HEPES pH 7.5, 150 mM NaCl, 0.5 mM TCEP.

### In vitro 20S degradation assay

HEK293T cells were transfected with HA-PPM1D or HA–BirA-K_0_ plasmids. Forty-eight hours after transfection, HA-tagged proteins were immunoprecipitated using HA magnetic agarose beads (AlpaLifeBio, KTSM1335). After extensive washing, HA-tagged proteins were eluted with HA peptide (MedChemExpress, HY-P0239) in 20 mM HEPES buffer (pH 8.0). Eluted protein, or 1 µg of recombinant H2B protein (Active Motif, 31492) was incubated with 0.5 µg of purified human 20S proteasome (Enzo, BML-PW8720) at 37 °C for 3 hours. For proteasome inhibition condition, the 20S proteasome was pre-incubated with 20 µM MG132 at 37 °C for 20 minutes prior to addition of substrates. Reactions were terminated by addition of SDS loading buffer followed by boiling at 100 °C for 6 min. Reaction products were analyzed by immunoblotting.

### In vitro ubiquitylation

0.2 µM UBA1, 1 µM UBE2D3, 0.2 µM UBR5, and 0.5 µM tr-PPM1D, which was generated as previously described, were mixed in the presence of 30 mM HEPES pH 7.5, 200 mM NaCl, 5mM MgCl_2_, and 4 mM ATP.^2^ Reactions were initiated by the addition of 50 µM ubiquitin. Reactions were conducted for the indicated time points at 37 °C and quenched with loading buffer. Reaction products were analyzed by immunoblotting using an anti-PPM1D antibody (Cell Signaling Technology, 11901) and Goat anti-Rabbit secondary (IRdye800CW Goat anti-rabbit, LiCor). Blots were imaged on a LI-COR CLx.

### Reporter-based PPM1D stability assay

MOLM13 cells were transduced with EGFP-PPM1D-IRES-mCherry virus. Forty-eight hours after infection, cells were selected with 1 µg/mL puromycin for 5 days. Ten days post-infection, cells were pre-treated with DMSO, 5 µM MG132, or 1 µM TAK-243 for 1 hour, followed by treatment with or without 50 µg/mL cycloheximide for 6 hours. Cells were then analyzed by flow cytometry using a BD FACSymphony A1 cell analyzer. Relative PPM1D protein levels were quantified as the ratio of GFP to mCherry mean fluorescence intensity (MFI).

### CRISPR screen for PPM1D degradation regulators

Ten percent (v/v) of the human genome-wide CRISPR-KO Brunello library was transduced to U937-Cas9 cells expressing EGFP-fl-PPM1D-IRES-mCherry, or EGFP-PPM1D_400-605_-IRES-mCherry. Twenty-four hours after infection, sgRNA-transduced cells were selected with 1 µg/mL puromycin for 5 days. Ten days after infection, cells were treated either DMSO (n = 3), or 50 µg/mL cycloheximide (n = 3) for 6 hours, then cell populations were separated using fluorescence-activated cell sorting (FACS). Four populations were collected (top 5%, top 5-15%, lowest 5%, and lowest 5-15%) based on GFP to mCherry mean fluorescent intensity (MFI) ratio on a BD FACSAria 1 cell sorter. Sorted cells were collected by centrifugation and subjected to direct lysis buffer reactions (1 mM CaCl_2_, 3 mM MgCl_2_, 1 mM EDTA, 1% Triton X-100, 10 mM Tris-HCl pH 7.5, supplemented with 0.2 mg/mL Proteinase K). The sgRNA sequence was amplified in a first PCR reaction with staggered forward primers. Twenty-five microliters of direct lysed cells were mixed with 2U Titanium Taq (Takara Bio, 639210), 0.5× Titanium Taq Buffer, dNTP mix, 1 µM forward stagger primers, and 1 µM common reverse primer in a 50 µL reaction. PCR program as follows: 94 °C 5 min, 27 cycles (94 °C 30 sec, 58 °C 15 sec, 72 °C 30 sec), 72 °C 2min. Two microliters of the primary PCR product were used as the template for the secondary PCR, where Illumina adapters and barcodes were added. After gel analysis, an equal amount of all samples was pooled and subjected to preparative agarose electrophoresis followed by gel purification. Sequencing was performed on a NovaSeq SP platform at the Broad Institute Clinical Labs.

The human BISON CRISPR-KO library, which contains 2,852 guide RNAs, targets 713 E1, E2, E3, deubiquitinases, and control genes.^3^ It was cloned into the pXPR003 by the genome perturbation platform (GPP, Broad Institute) as previously described. U937-Cas9 expressing EGFP-fl-PPM1D-IRES-mCherry or EGFP-tr-PPM1D-IRES-mCherry were transduced with the library virus, cells were selected with 1 µg/mL puromycin and collected ten days after infection, then were processed as described above.

### Alkaline Comet assay

Comet assays were conducted as previously described.^4,5^ Cells were resuspended to 1×10^5^ cells/mL and mixed with 1% low-melting agarose (R&D Systems, 4250-500-02) at a 1:10 ratio and plated on comet slides (R&D Systems, 4250-200-03). Cells were then lysed and immersed in alkaline unwinding solution as per manufacturer’s instruction. Fluorescence microscopy was performed at 10X magnification using a Nikon Eclipse Ti inverted microscope and analysis of comet tails were performed using the OpenComet software.^6^ At least 160 comet tails were measured per condition.

### HR and NHEJ reporter assay

HR and NHEJ rates were assessed using a CRISPR/Cas9 engineered U2OS cell line expressing fl-PPM1D-only and a U2OS parental cell line with integrated HR (DR-GFP) or NHEJ (EJ5-GFP) reporter construct essentially as described.^7^ Briefly, two days after transfection of siRNAs, cells were re-seeded to a 24-well plate. The next day, the cells were transfected with I-SceI plasmid (Addgene, 60960). Cells were collected 48 hours after transfection and subjected to flow cytometry analysis to examine the proportion of GFP-positive cells.

### Multicolor competitive assay

The multicolor competition assay was performed as previously described.^8^ U2OS parental and isogenic cell lines were transduced with lentiviruses expressing either RFP or BFP and selected with 5 µg/mL puromycin. RFP-labeled cells were transfected with *UBR5* siRNA, whereas BFP-labeled cells were transfected with non-targeting control siRNA. Forty-eight hours after transfection, equal numbers of RFP and BFP cells were mixed, and 24 hours later the mixed populations were exposed to 2.5 Gy irradiation. Cells were analyzed by flow cytometry 72 hours after irradiation. Relative survival was calculated as the ratio of RFP-positive to BFP-positive cells. All data were normalized to control mixtures in which both RFP- and BFP-labeled cells were transfected with non-targeting control siRNA and processed in parallel.

### Proximity-dependent Biotin Identification assay (BioID)

HEK293T cells were transduced with lentiviruses expressing BirA-fl-PPM1D or BirA-tr-PPM1D. Forty-eight hours after infection, cells were selected with puromycin for 5 days. Ten days post-infection, cells were seeded into 6-cm dishes at approximately 70% confluency. The following day, cells were treated with 50 µM biotin for 24 hours. Six hours before harvest, cells were treated with 10 µM MG132 in the presence or absence of 50 µg/mL cycloheximide. Cells were harvested in NP40 buffer supplemented with 10 µM MG132 and 1× Halt Protease Inhibitor Cocktail. Lysates were probe-sonicated and cleared by centrifugation at 14,000 rpm for 20 min at 4 °C. For each sample, 100 µL of pre-cleared streptavidin Sepharose beads (Cytiva, 17511301) were added, and samples were incubated for 3 hours at 4 °C with end-to-end rotation. Following incubation, beads were washed three times with SDS wash buffer (2% SDS, 50 mM Tris-HCl, pH 7.5), three times with BioID wash buffer (500 mM NaCl, 0.4% SDS, 50 mM Tris-HCl, pH 7.5), and six times with 50 mM Tris-HCl (pH 7.5). After the final wash, beads were resuspended in 50 mM Tris-HCl (pH 7.5) for mass spectrometry or immunoblotting analysis. Samples were analyzed by mass spectrometry at the Taplin Mass Spectrometry Facility at Harvard Medical School. Briefly, gel slices from Coomassie blue-stained SDS-PAGE were cut into approximately 1 mm3 pieces, and subjected to trypsin digestion. Peptides were extracted and reconstituted in 5–10 µl of HPLC solvent A (2.5% acetonitrile, 0.1% formic acid). A nano-scale reverse-phase HPLC capillary column was created by packing 2.6 µm C18 spherical silica beads into a fused silica capillary. After equilibrating the column each sample was loaded via a Famos auto sampler (LC Packings, San Francisco, CA) onto the column. A gradient was formed, and peptides were eluted with increasing concentrations of solvent B (97.5% acetonitrile, 0.1% formic acid). As peptides eluted, they were subjected to electrospray ionization and then entered an LTQ Orbitrap Velos Pro ion-trap mass spectrometer (Thermo Fisher Scientific). Peptides were detected, isolated and fragmented to produce a tandem mass spectrum of specific fragment ions for each peptide. Raw data were searched against a human protein database (FASTA, January 6, 2023) using the Sequest search algorithm. Peptide-spectrum matches were filtered using linear discriminant analysis to achieve an estimated false discovery rate of approximately 1%.

### Depmap and TCGA analysis

For **Figures 1A-B** and **Supplemental Figure 1A**, *PPM1D* truncating mutations, copy number gain, and expression data was obtained from the Catalogue of Somatic Mutations in Cancer (COSMIC) database (https://cancer.sanger.ac.uk/cosmic, v103).^9^ For **Figures 5J-K**, data for The Cancer Genome Atlas (TCGA) studies was obtained from cBioportal (www.cBioportal.org) and data from the Cancer Dependency Map was obtained from Depmap (www.depmap.org). *UBR5* mutations were restricted to truncating (nonsense or frameshift) to maximize likelihood of pathogenicity. *UBR5* and *PPM1D* amplifications from TCGA were defined as previously described.^10^ All data was obtained from the aforementioned sources in February 2026.

## QUANTIFICATION AND STATISTICAL ANALYSIS

All quantifications are presented as the mean ± SEM unless indicated. Significance was determined by unpaired two-tailed Student’s t-test. All RT-qPCR quantifications are presented as the mean of three technical repeats from three independent biological replicates. ImageJ software was used for quantification and statistics of protein bands from scanned blots for degradation assays. Statistical analyses were performed by using GraphPad Prism 10.

