## Supplemental figures for "Distinct Proteasomal Pathways Drive Oncogenic PPM1D Activation"

### Figure S1

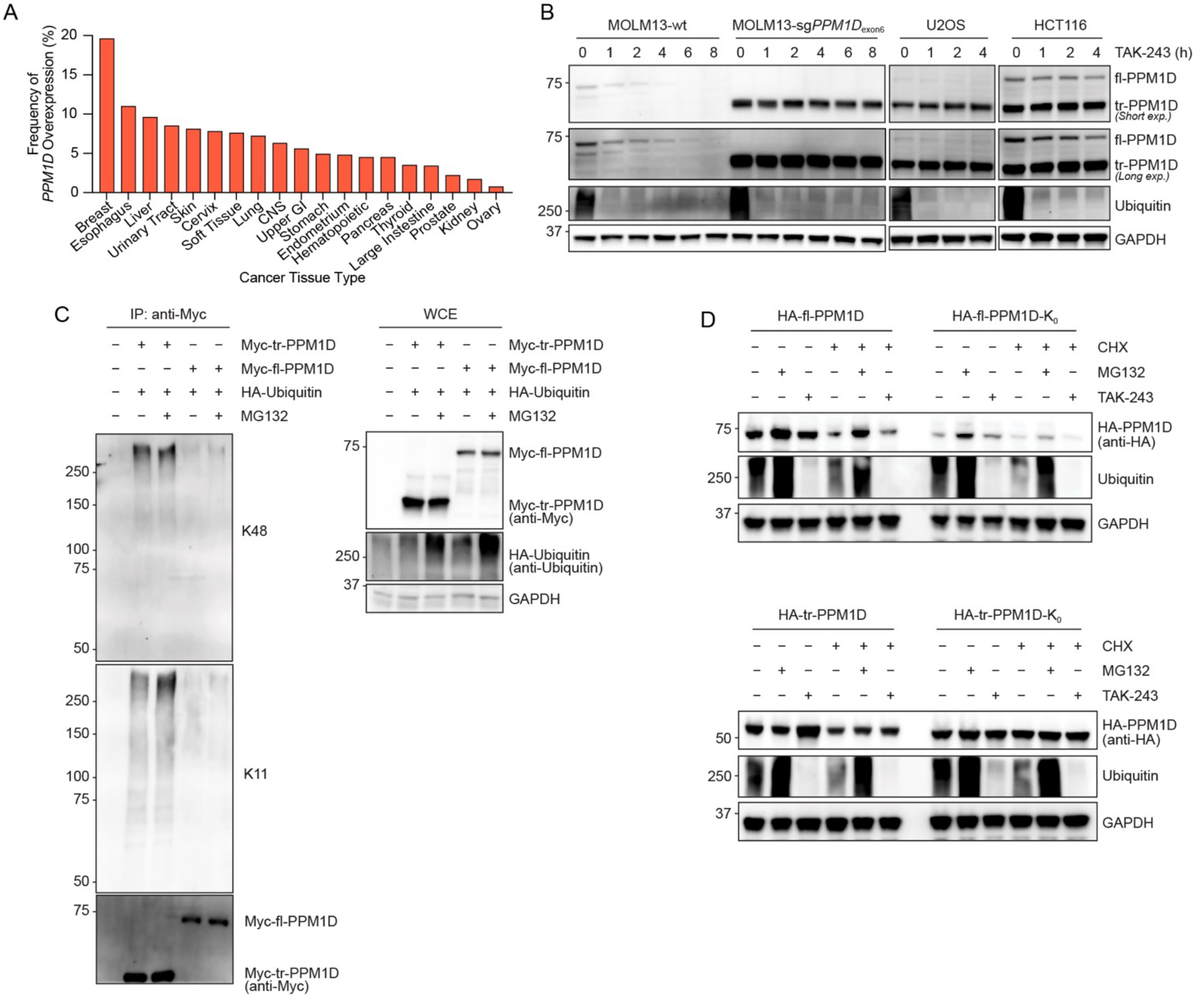


**Figure S1. fl-PPM1D and tr-PPM1D undergo distinct modes of proteasomal degradation.**

(A) Histogram showing the frequency of *PPM1D* overexpression by cancer tissue type acquired from the Catalogue of Somatic Mutations in Cancer (COSMIC) database.

(B) Western blot time-course of PPM1D in in MOLM13-wt, MOLM13-sg*PPM1D*$_{exon6}$, U2OS, or HCT116 after treatment with TAK-243.

(C) Myc-tr-PPM1D or Myc-fl-PPM1D were co-expressed with HA-Ubiquitin in HEK293T treated with DMSO or MG132 cells followed by denaturing immunoprecipitation of Myc-PPM1D and immunoblotting for Ubiquitin-K11 and Ubiqiutin-K48. Whole cell extract (WCE) is shown on the right.

(D) Western blots of HA-PPM1D in HEK293T cells expressing HA-fl-PPM1D, HA-fl-PPM1D-$K_0$, HA-tr-PPM1D, or HA-tr-PPM1D-$K_0$ after treatment with cycloheximide (CHX), MG132, or TAK-243.

### Figure S2

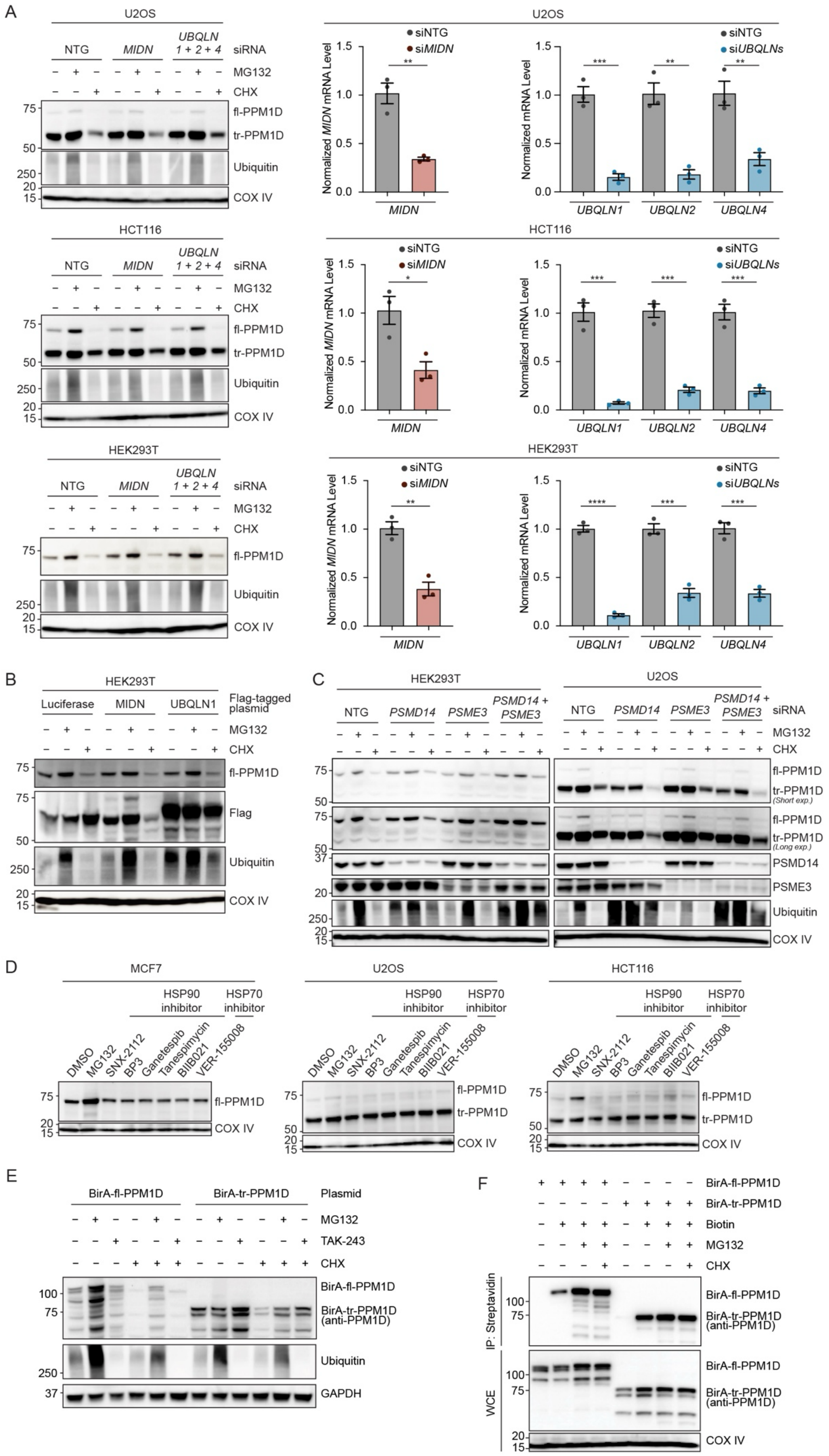

A
U2OS
NTG
MIDN
UBQLN 1 + 2 + 4
siRNA
MG132
CHX
fl-PPM1D
tr-PPM1D
Ubiquitin
COX IV
siNTG
siMIDN
siUBQLNs
Normalized MIDN mRNA Level
Normalized mRNA Level
UBQLN1
UBQLN2
UBQLN4
HCT116
HEK293T
B
Luciferase
UBQLN1
Flag-tagged plasmid
Flag
C
PSMD14
PSME3
PSMD14 + PSME3
(Short exp.)
(Long exp.)
D
MCF7
HSP90 inhibitor
HSP70 inhibitor
DMSO
SNX-2112
BP3
Ganetespib
Tanespimycin
BIIB021
VER-155008
E
BirA-fl-PPM1D
BirA-tr-PPM1D
Plasmid
TAK-243
BirA-tr-PPM1D (anti-PPM1D)
GAPDH
F
Biotin
IP: Streptavidin
WCE

**Figure S2. Ubiquitin-independent proteasomal degradation of fl-PPM1D is not regulated by known chaperones.**

(A) Western blot (left) of PPM1D following siRNA mediated knockdown of non-targeting control (NTG), *MIDN*, or *UBQLN1/2/4* and treatment with MG132 or cycloheximide (CHX) in U2OS (top), HCT116 (middle), and HEK293T (bottom) cells. Normalized mRNA levels (right) of *MIDN* and *UBQLN1/2/4* after NTG, *MIDN*, or *UBQLN1/2/4* silencing were determined by quantitative PCR. Data are presented as mean ± SEM (n = 3 biological replicates). Statistical significance was determined using unpaired two-tailed Student's t-test; *$p<0.05$, **$p<0.01$, ***$p<0.001$, ****$p<0.0001$.

(B) Western blot of PPM1D following overexpression of FLAG-tagged luciferase control, MIDN, or UBQLN1 and treatment with MG132 or CHX in HEK293T cells.

(C) Western blot of PPM1D, PSMD14, and PSME3 following siRNA mediated knockdown of NTG, *PSMD14*, *PSME3*, or *PSMD14* and *PSME3* and treatment with MG132 or CHX in HEK293T (left) and U2OS (right) cells.

(D) Western blot of PPM1D following treatment of DMSO, MG132, HSP90 inhibitors, or HSP70 inhibitors in MCF7, U2OS, or HCT116 cells.

(E) Western blot of BirA-PPM1D in HEK293T cells expressing BirA-fl-PPM1D or BirA-tr-PPM1D and treated with MG132, TAK-243, or CHX.

(F) HEK293T cells expressing BirA-fl-PPM1D or BirA-tr-PPM1D were incubated with biotin for 24 hours, followed by MG132 or MG132 and CHX treatment. Biotinylated substrates were enriched by streptavidin and analyzed by immunoblotting for PPM1D. Whole cell extract (WCE) is shown below.

#### Figure S3

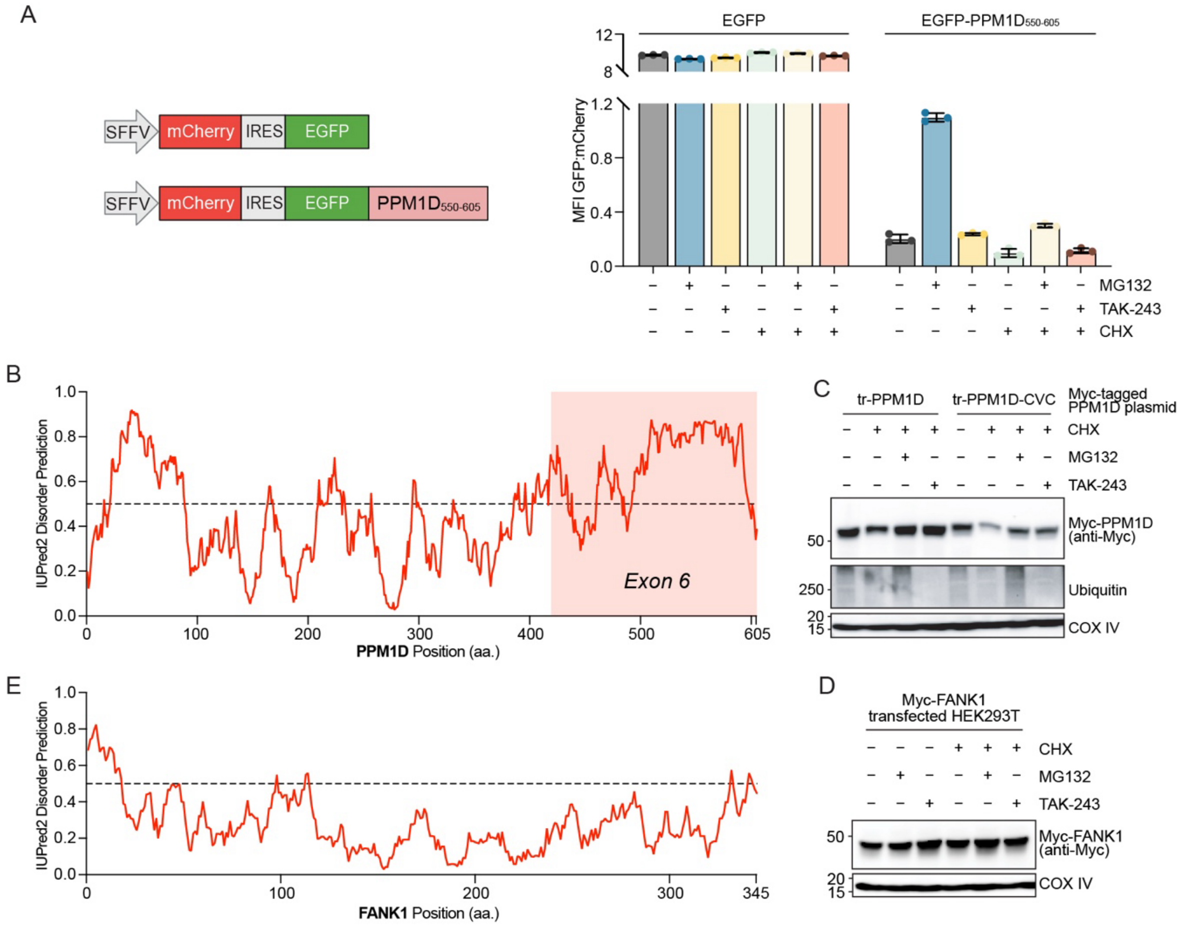


**Figure S3. Ubiquitin-independent proteasomal degradation of fl-PPM1D is mediated by the C-terminus.**

(A) Schematic (left) and GFP:mCherry quantification by flow cytometry (right) in MOLM13 cells expressing the EGFP-IRES-mCherry or EGFP-PPM1D$_{500\text{-}605}$-IRES-mCherry stability reporter treated with CHX, MG132, or TAK-243. Data are presented as mean ± SD (n = 3 biological replicates).

(B) Disorder tendency of fl-PPM1D was predicted using IUPred2 (https://iupred2a.elte.hu/). Scores above 0.5 indicate regions with a high propensity for intrinsic disorder.

(C) Western blot of Myc-PPM1D in HEK293T cells expressing Myc-tr-PPM1D or Myc-tr-PPM1D-CVC and treated with CHX, MG132, or TAK-243.

(D) Western blot of Myc-FANK1 in HEK293T cells expressing Myc-FANK1 and treated with CHX, MG132, or TAK-243.

(E) Disorder tendency of FANK1 was predicted using IUPred2. Scores above 0.5 indicate regions with a high propensity for intrinsic disorder.

### Figure S4

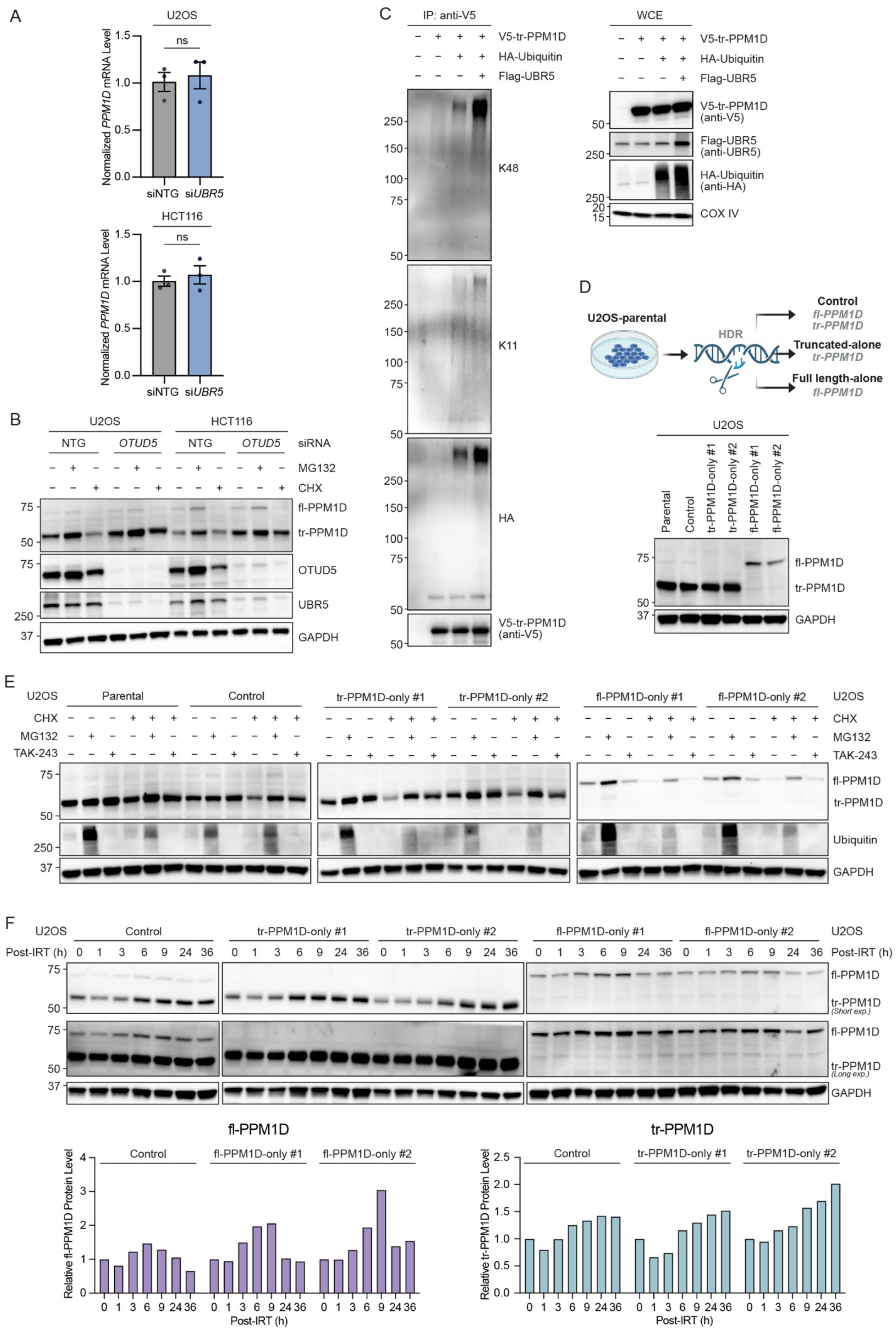
A
U2OS
HCT116
Normalized PPM1D mRNA Level
ns
siNTG
siUBR5
B
C
IP: anti-V5
WCE
V5-tr-PPM1D
HA-Ubiquitin
Flag-UBR5
K48
K11
HA
D
U2OS-parental
HDR
Control
Truncated-alone
Full length-alone
fl-PPM1D
tr-PPM1D
GAPDH
E
F
Post-IRT (h)
Relative fl-PPM1D Protein Level
Relative tr-PPM1D Protein Level

**Figure S4. UBR5 regulates the ubiquitin-dependent proteasomal degradation of tr-PPM1D.**

(A) Normalized mRNA levels of *PPM1D* after siRNA mediated knockdown of non-targeting control (NTG) or *UBR5* in U2OS (top) or HCT116 (bottom) determined by quantitative PCR. Data are presented as mean ± SEM (n = 3 biological replicates). Statistical significance was determined using unpaired two-tailed Student's t-test; ns indicates non-significant ($p$>0.05).

(B) Western blot of PPM1D, OTUD5, and UBR5 following siRNA mediated knockdown of non-targeting control (NTG) or *OTUD5* and treatment with MG132 or cycloheximide (CHX) in U2OS and HCT116 cells.

(C) V5-tr-PPM1D, HA-Ubiquitin, and FLAG-UBR5 were overexpressed in HEK293T cells followed by denaturing immunoprecipitation of V5-tr-PPM1D and immunoblotting for Ubiquitin-K11, Ubiqiutin-K48, and HA-Ubiquitin. Whole cell extract (WCE) is shown the on right.

(D) Schematic of the process of generating U2OS-control, tr-PPM1D-only, or fl-PPM1D-only isogenic cell lines using CRISPR/Cas9-mediated Homology-Directed Repair (HDR) (top), and western blot of PPM1D in the isogenic cell lines (bottom).

(E) Western blot of PPM1D in U2OS isogenic cell lines after treatment with CHX, MG132, or TAK-243.

(F) Western blot time-course (top) and quantification (bottom) of fl- or tr-PPM1D in U2OS isogenic cell lines after exposure to 2.5 Gy irradiation.

### Figure S5

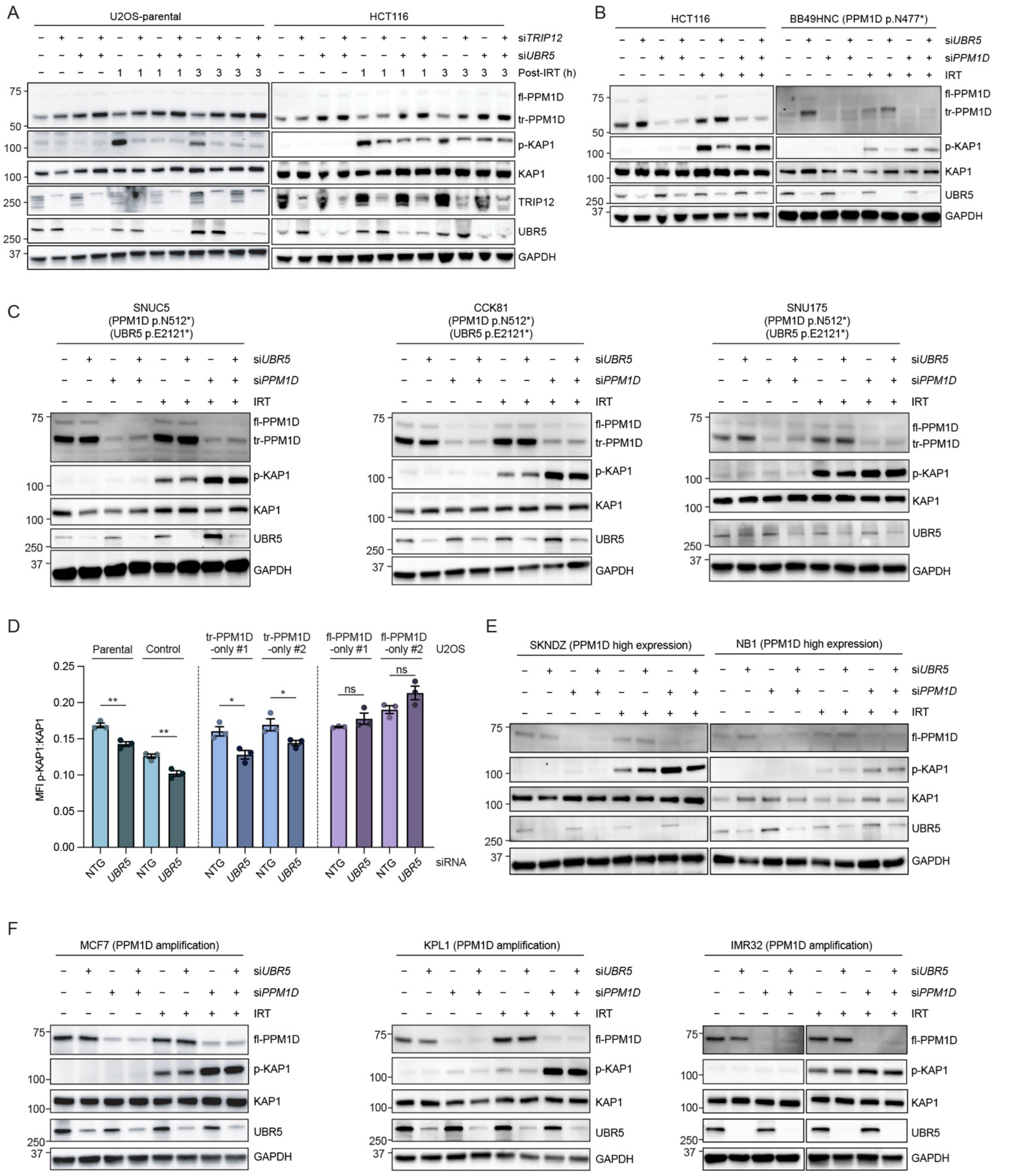


**Figure S5. UBR5 regulation of p-KAP1 is mediated by truncated PPM1D.**

(A) Western blot time-course of PPM1D, p-KAP1, KAP1, TRIP12, and UBR5 in U2OS and HCT116 cells after siRNA mediated knockdown of non-targeting control (NTG), *TRIP12*, *UBR5,* or *TRIP12* and *UBR5* and exposure to 0 or 2.5 Gy irradiation.

(B) Western blot of PPM1D, p-KAP1, KAP1, and UBR5 in HCT116 and BB49HNC cells after siRNA mediated knockdown of NTG, *UBR5, PPM1D,* or *UBR5* and *PPM1D* and exposure to 0 or 2.5 Gy irradiation.

(C) Western blot of PPM1D, p-KAP1, and KAP1 in SNUC5, CCK81, SNU175 cells (all with *PPM1D* truncating mutations and loss-of-function *UBR5* mutations) after siRNA mediated knockdown of NTG or *UBR5* and exposure to 0 or 2.5 Gy irradiation.

(D) p-KAP1:KAP1 mean fluorescence intensity (MFI) quantified by intracellular flow cytometry in U2OS parental or isogenic cell lines after siRNA mediated knockdown of NTG or UBR5 and treatment with 2.5 Gy irradiation. Data are presented as mean ± SEM (n = 3 biological replicates). Statistical significance was determined using unpaired two-tailed Student's t-test; ns indicates non-significant ($p>0.05$), ***$p<0.001$, ****$p<0.0001$.

(E) Western blot of PPM1D, p-KAP1, KAP1, and UBR5 in SKNDZ and NB1 cells (all with wild-type *PPM1D* high expression) after siRNA mediated knockdown of NTG or *UBR5* and exposure to 0 or 2.5 Gy irradiation.

(F) Western blot of PPM1D, p-KAP1, KAP1, and UBR5 in MCF7, KPL1, and IMR32 cells (all with wild-type *PPM1D* amplification) after siRNA mediated knockdown of NTG or *UBR5* and exposure to 0 or 2.5 Gy irradiation.

#### Figure S6

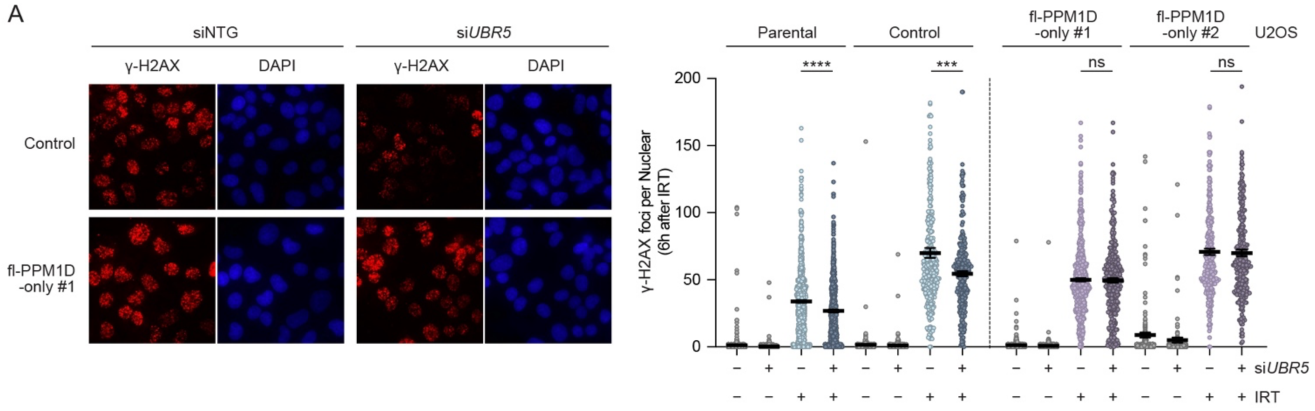


**Figure S6. UBR5 regulation of γ-H2AX foci formation is mediated by truncated PPM1D.**

(A) Immunofluorescence of γ-H2AX was performed in U2OS parental and isogenic cell lines transfected with non-targeting control (NTG) or *UBR5* siRNA, 6 hours after 2.5 Gy irradiation. Representative images are shown on the left, and quantification of γ-H2AX foci per nucleus is shown on the right. n>190 cells for each condition were analyzed using CellProfiler. Data are presented as mean ± SEM. Statistical significance was determined using unpaired two-tailed Student's t-test; ns indicates non-significant ($p$>0.05), *$p$<0.05, **$p$<0.01.
